# HEATR5B associates with dynein-dynactin and selectively promotes motility of AP1-bound endosomal membranes

**DOI:** 10.1101/2023.03.14.532574

**Authors:** Vanesa Madan, Lucas Albacete Albacete, Li Jin, Pietro Scaturro, Joseph L. Watson, Nadine Muschalik, Farida Begum, Jérôme Boulanger, Karl Bauer, Michael A. Kiebler, Emmanuel Derivery, Simon L. Bullock

**Author notes:** Joint second authors (Lucas Albacete Albacete, Li Jin). Department of Biochemistry, University of Washington, Seattle, WA, USA.

## Abstract

The dynein motor complex mediates polarised trafficking of a wide variety of organelles, intracellular vesicles and macromolecules. These functions are dependent on the dynactin complex, which helps recruit cargoes to dynein’s tail region and activates motor movement. How dynein and dynactin orchestrate trafficking of diverse cargoes is unclear. Here, we identify HEATR5B, an interactor of the AP1 clathrin adaptor complex, as a novel player in dynein-dynactin function. HEATR5B is one of several proteins recovered in a biochemical screen for proteins whose association with the human dynein tail complex is augmented by dynactin. We show that HEATR5B binds directly to the dynein tail and dynactin and stimulates motility of AP1-associated endosomal membranes in human cells. We also demonstrate that the HEATR5B homologue in *Drosophila* is an essential gene that promotes dynein-based transport of AP1-bound membranes to the Golgi apparatus. As HEATR5B lacks the coiled-coil architecture typical of dynein adaptors, our data point to a non-canonical process orchestrating motor function on a specific cargo. We additionally show that HEATR5B promotes association of AP1 with endosomal membranes in a dynein-independent manner. Thus, HEATR5B co-ordinates multiple events in AP1-based trafficking.

## INTRODUCTION

Microtubule motors play a central role in the trafficking of cellular constituents through the cytoplasm. Whilst multiple kinesin family members are tasked with transporting cargoes towards microtubule plus ends, a single motor – cytoplasmic dynein-1 (dynein) – is responsible for almost all minus end-directed movement (Reck-Peterson *et al*., 2018). Dynein’s diverse cellular cargoes include mRNAs, protein complexes, nuclei, mitochondria, lysosomes, the Golgi apparatus and multiple classes of vesicle. It is unclear how one motor orchestrates trafficking of so many cargoes.

Dynein is a highly conserved, 1.3-MDa complex of six subunits, which are each present in two copies (Reck-Peterson *et al*., 2018; Schmidt & Carter, 2016). The heavy chain subunit contains a motor domain and a tail domain (Figure 1A). The motor domain has force-generating ATPase activity and a microtubule-binding site, which work in concert to drive movement along the track. The tail domain mediates homodimerisation and recruits the other dynein subunits, which make important contributions to complex stability and cargo binding (Lee *et al*., 2018; Reck-Peterson *et al*., 2018; Schroeder *et al*., 2014).

**Figure 1.**
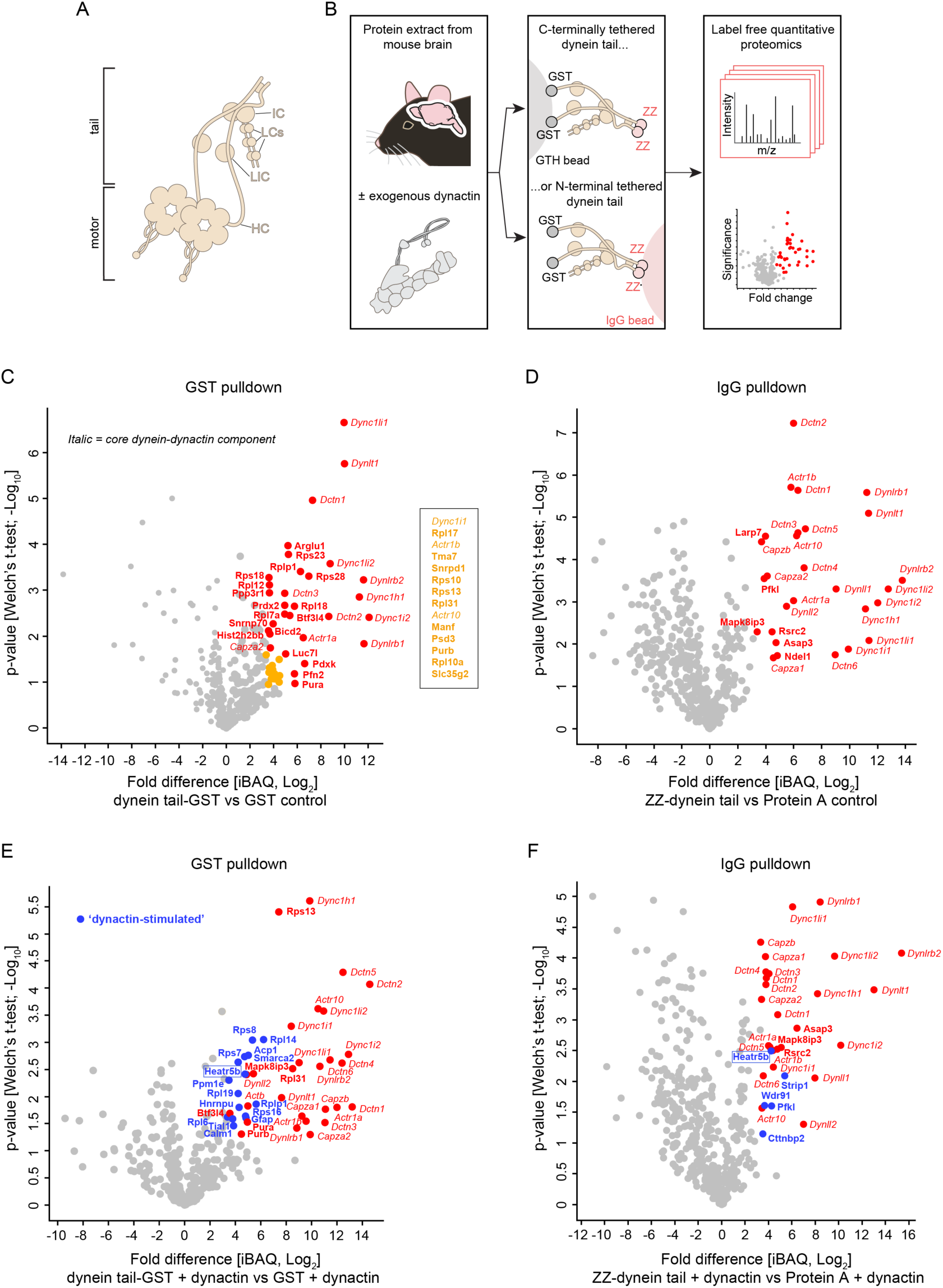
Dynein architecture and identification of novel dynein tail interactors. (A) Overview of dynein complex. IC, intermediate chain; LCs, light chains; LIC, light intermediate chain; IC, intermediate chain. (B) Biochemical strategy for identifying dynein tail interactors, including those whose association is stimulated by dynactin. ZZ is a protein A-based tag. (C – E) Volcano plots showing results of label-free quantitative proteomics (iBAQ, intensity based absolute quantification). Proteins that met our criteria for enrichment on the dynein tail vs the control (median log_2_(fold change) >3.322 and Welch’s t-test q-value <0.05) are labelled in colour and named. For clarity, the names of significant hits that cluster closely together on the plot (yellow circles) are given separately. Core dynein-dynactin components that were enriched on the dynein tail are shown in italics, with other enriched proteins shown in bold; of these ‘none core’ proteins, those only detected in the presence of exogenous dynactin are shown in bold and blue.

*In vitro* studies have shown that mammalian dynein needs additional factors for processive movement on microtubules (McKenney *et al*., 2014; Miura *et al*., 2010; Schlager *et al*., 2014; Trokter *et al*., 2012). The paradigmatic activation mechanism uses a combination of the 1.1-MDa dynactin complex and one of a number of coiled coil-containing proteins – so called ‘activating adaptors’ (Olenick & Holzbaur, 2019; Reck-Peterson *et al*., 2018) – that interact with cargo-associated proteins. The activating adaptor stimulates the interaction of dynein’s tail with dynactin, which re-positions the motor domains for processive movement (McKenney *et al*., 2014; Schlager *et al*., 2014; Splinter *et al*., 2012; Zhang *et al*., 2017). Dynactin is also important for cargo recruitment to dynein as it stabilises the association of the motor with activating adaptors (McKenney *et al*., 2014; Schlager *et al*., 2014; Splinter *et al*., 2012).

The activating adaptors and cargo-associated proteins that connect different cargoes to dynein’s tail and dynactin have been defined in only a small number of cases (Hoogenraad & Akhmanova, 2016; Olenick & Holzbaur, 2019; Reck-Peterson *et al*., 2018). Thus, for many cargoes the proteins that provide the bridge to the motor complex and activate motor movement are not known. Identifying such factors is a prerequisite for understanding principles of cargo recognition and motor activation, as well as for dissecting the cellular functions of specific dynein-based transport processes.

We set out to address this issue by identifying novel biochemical interactors of the dynein tail, including those whose binding is enhanced by dynactin. Functional analysis of one dynactin-stimulated interactor, the non-coiled-coil protein HEATR5B, shows that it binds directly to the dynein tail and dynactin and has an evolutionarily conserved function in promoting motility of AP1-associated endosomal membranes. This work reveals a critical contribution of a protein that lacks coiled-coil architecture to dynein-based transport and provides novel insights into dynein functions during membrane trafficking. We also show that HEATR5B promotes association of AP1 with endosomal membranes independently of dynein. Thus, we have identified a factor that co-ordinates the recruitment of AP1 to endosomal membranes with microtubule-based transport of these structures.

## RESULTS

### A biochemical screen for dynein tail interactors

It was recently found that autoinhibitory interactions involving the dynein motor domain reduce association of the dynein tail with dynactin and cargo adaptors (Htet *et al*., 2020; Zhang *et al*., 2017). We therefore sought to increase the likelihood of identifying novel tail interactors by performing affinity purifications with a recombinant human dynein complex that lacks the motor domains (Figure 1B and Figure S1). This ‘tail complex’ (comprising residues 1-1079 of the DYNC1H1 heavy chain, the DYNC1I2 intermediate chain, the DYNC1LI2 light intermediate chain and the DYNLL1, DYNRB1 and DYNLT1 light chains) was produced in insect cells and coupled to beads via epitope tags. The beads were then incubated with extracts from mouse brain, which provides a concentrated source of potential binding partners. Pull-downs were performed with both N-terminally and C-terminally tethered tail complexes to prevent loss of any interactions obstructed by coupling to beads in a specific orientation. In a subset of pull-downs, brain extracts were spiked with purified dynactin (Figure S1), which we reasoned would facilitate capture of tail interactors involved in cargo transport processes. Following mild washing of beads, retained proteins from three technical replicates per condition were analysed with label-free quantitative proteomics (Supplementary Datasets S1-S4).

In the absence of exogenous dynactin, 57 proteins met our criteria for enrichment in the dynein tail pull-downs compared to controls in which only the epitope tags were coupled to the beads (log_2_(fold change) >3.322 (i.e. (fold change) >10) and Welch’s t-test q-value <0.05; Figure 1C, D). 22 of these proteins were core components of dynein or dynactin (Figure 1C, D – italics). These factors included isoforms of dynein subunits that were not present in the recombinant human tail complex (Dync1i1, Dync1li1, Dynll2 and Dynlrb2), which were presumably recovered either because they exchange with their counterparts in the recombinant tail complex or are part of a second dynein complex that can be bridged by dynactin (Grotjahn *et al*., 2018; Urnavicius *et al*., 2018). The other 35 proteins specifically captured by the tail (Figure 1C, D – bold; Table S1) included two known coiled-coil containing cargo adaptors for dynein-dynactin – BicD2 and Mapk8ip3 (also known as Jip3) – and the dynein regulator Ndel1 (Reck-Peterson *et al*., 2018). The other tail-enriched factors had not previously been shown to interact with dynein or dynactin. These proteins included ribosomal subunits, RNA binding proteins and other proteins with diverse biochemical functions (Table S1).

We next determined which proteins were captured on the dynein tail versus the epitope tag controls when exogenous dynactin was spiked into the extracts. In addition to dynein and dynactin components (Figure 1E, F – italics), 28 proteins were enriched on the tail with dynactin spiking (Figure 1E, F – bold). Seventeen of these proteins were not captured by the tail in the absence of exogenous dynactin (Figure 1E, F – bold and blue; Table S2). These ‘dynactin-stimulated’ interactors included Strip1 (Striatin-interacting protein 1), a component of the Stripak (Striatin-interacting phosphatases and kinases) complex, and the Stripak-associated factor Cttnbp2 (Cortactin binding protein 2) (Kuck *et al*., 2019). *Drosophila* Stripak components associate with dynein and dynactin and regulate transport of endosomes, autophagosomes and dense-core vesicles (Neisch *et al*., 2017; Sakuma *et al*., 2014). Our data strengthen evidence that interactions of STRIPAK proteins with dynein-dynactin are conserved in mammals (Goudreault *et al*., 2009). The other dynactin-stimulated proteins had not been linked with dynein or dynactin in previous studies.

Collectively, our pull-down experiments identified ∼50 novel interacting proteins of the dynein tail, several of which had their association enhanced by dynactin.

### HEATR5B associates with the dynein tail and dynactin

From our list of candidate tail interactors, we were particularly drawn to Heatr5B (Heat repeat containing protein 5B; also known as p200a (Hirst *et al*., 2005)) because this protein was the only factor whose recruitment to both the N-terminally and C-terminally tethered dynein tail complexes was stimulated by exogenous dynactin (boxed labels in Figure 1E, F). Moreover, a previous proteomic study found that Heatr5B is present on dynactin-associated membranes isolated from mouse brain (Hinckelmann *et al*., 2016).

Heatr5B (HEATR5B in humans) is a 225 kDa protein that lacks a coiled-coil domain and is predicted to mostly comprise HEAT repeats (Yoshimura & Hirano, 2016); https://www.uniprot.org/uniprot/Q9P2D3). Humans also have a related protein, HEATR5A (p200b), which was not recovered in our screen for tail interactors. A complex of HEATR5B, Aftiphilin (AFTPH/AFTIN) and γ-synergin (SYNRG/AP1GBP1) interacts with the AP1 (Adaptor protein-1 (AP1) complex and the Golgi-localized, gamma adaptin ear-containing, ARF-binding (GGA) protein (Hirst *et al*., 2005; Lui *et al*., 2003), adaptors that orchestrate formation and cargo loading of a subset of clathrin-coated vesicles from intracellular membranes (Nakatsu *et al*., 2014; Nakayama & Wakatsuki, 2003; Sanger *et al*., 2019). Neither AFTPH or SYNRG were found in our dynein tail pulldowns, suggesting they do not associate with the motor complex or interact less strongly with it than HEATR5B does. RNAi-mediated knockdowns have implicated HEATR5B in AP1-mediated cargo sorting in HeLa cells (Hirst *et al*., 2005), *Drosophila* imaginal discs (Le Bras *et al*., 2012) and *C. elegans* epidermal cells (Gillard *et al*., 2015). The importance of HEATR5B proteins is underscored by the recent finding that hypomorphic mutations in the human gene are associated with the neurodevelopmental syndrome pontocerebellar hypoplasia (Ghosh *et al*., 2021). However, the molecular function of HEATR5B is not clear in any of these systems.

We first asked if HEATR5B is part of a complex with dynein and dynactin in human cells by performing immunoprecipitations from HEK293 cell lines that stably express either GFP-tagged HEATR5B or GFP alone. As expected, association of AFTPH, SYNRG and the AP1γ subunit was detected with GFP-HEATR5B but not the GFP control (Figure 2A). Dynein and dynactin components were also specifically precipitated with GFP-HEATR5B (Figure 2A), corroborating the results of our tail pull-down experiments. To determine if HEATR5B can interact directly with dynein or dynactin, we performed *in vitro* pull-downs with purified proteins. Whilst truncated versions of HEATR5B were unstable and could not be purified, we could produce a full-length GFP-tagged version of the protein in insect cells (Figure S1). We detected binding of the recombinant dynein tail complex to recombinant GFP-HEATR5B but not GFP alone (Figure 2B). Purified dynactin also interacted specifically with recombinant GFP-HEATR5B (Figure 2B). Our previous finding that association of HEATR5B with the dynein tail in brain extracts is stimulated by dynactin (Figure 1E, F) suggests that HEATR5B can interact simultaneously with both complexes. Compatible with this notion, we did not observe competition between the purified dynein tail and dynactin for HEATR5B binding in our *in vitro* binding assay when both complexes were added simultaneously to the beads (Figure 2B). However, we cannot rule out the possibility that a competitive interaction was masked by binding sites on one of the components not being saturated. Nonetheless, we can conclude from this set of experiments that HEATR5B complexes with endogenous dynactin and dynein in cell extracts and can interact with both complexes directly.

**Figure 2.**
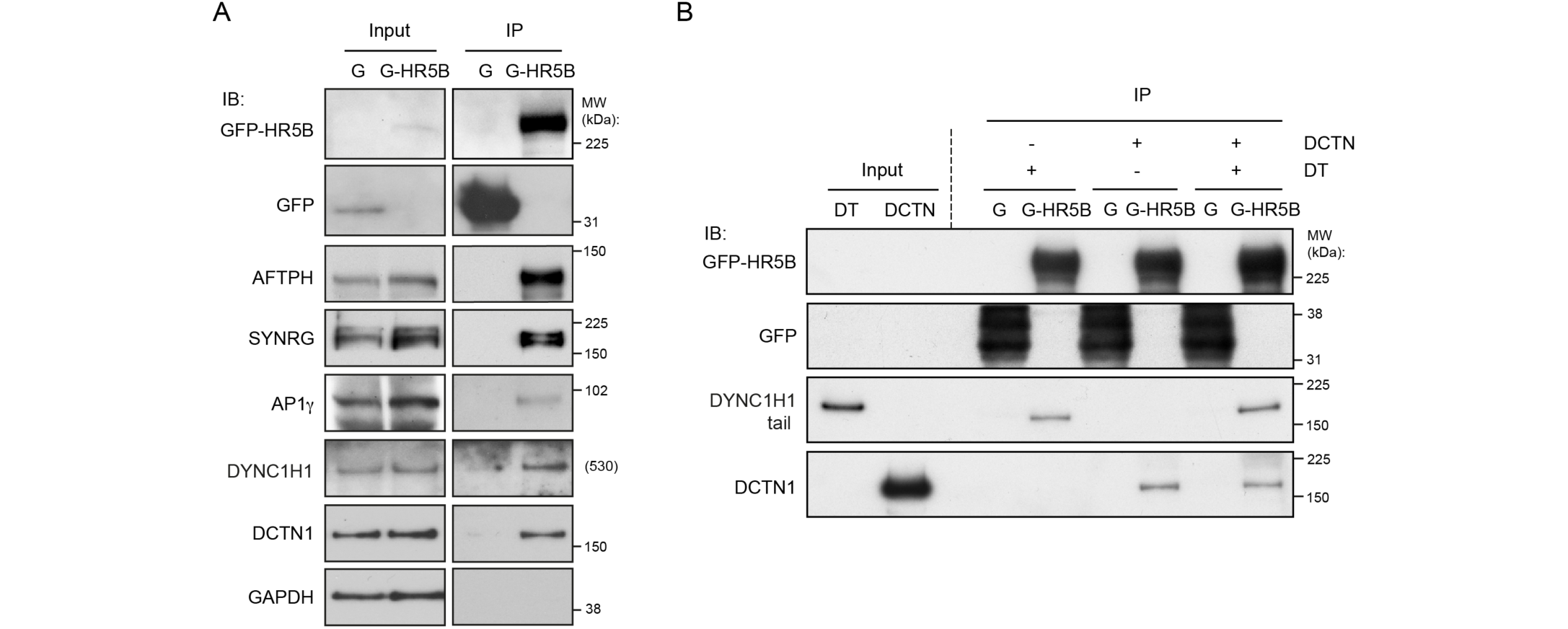
Confirmation of HEATR5B interaction with dynein and dynactin. **(A, B)** Immunoblots showing results of (A) immunoprecipitation experiments from GFP-HEATR5B and GFP HEK293 cells and (B) *in vitro* pull-downs with purified, bead-associated GFP-tagged human HEATR5B or GFP protein incubated with purified dynein tail and/or dynactin complexes. G-HR5B, GFP-HEATR5B; G, GFP; IP, immunoprecipitate; IB, protein probed by immunoblotting; MW, molecular weight of protein standards (theoretical MW is shown for DYNC1H1 as standards of a similar molecular weight were not available); DT, dynein tail complex; DCTN, dynactin complex. In (A), GAPDH is a negative control.

### HEATR5B is co-transported with AP1-positive endosomal membranes

We next investigated the possibility that HEATR5B is part of a link between AP1-positive membranes and dynein-dynactin. We first asked if HEATR5B co-localises with AP1. Although antibodies to HEATR5B work in immunoblots, they are not suitable for immunostaining of cells. We therefore stained HeLa cells that have a stable integration of the GFP-HEATR5B construct with an antibody to the γ subunit of the adaptor complex (AP1γ). GFP-HEATR5B was enriched in small puncta in the cytoplasm, ∼ 30% of which co-localised with AP1γ puncta (Figure 3A, B). In contrast, the population of AP1γ that associated with the perinuclear *trans*-Golgi network (TGN, marked with TGN46 antibodies) rarely overlapped with GFP-HEATR5B (Figure 3A and Figure S2A, B). GFP-HEATR5B puncta also seldomly associated with EEA1-positive early endosomes or LAMP1-positive lysosomal membranes (Figure 3B and Figure S2B). These results reveal that HEATR5B is selectively enriched on AP1-positive structures in the cytoplasm.

**Figure 3.**
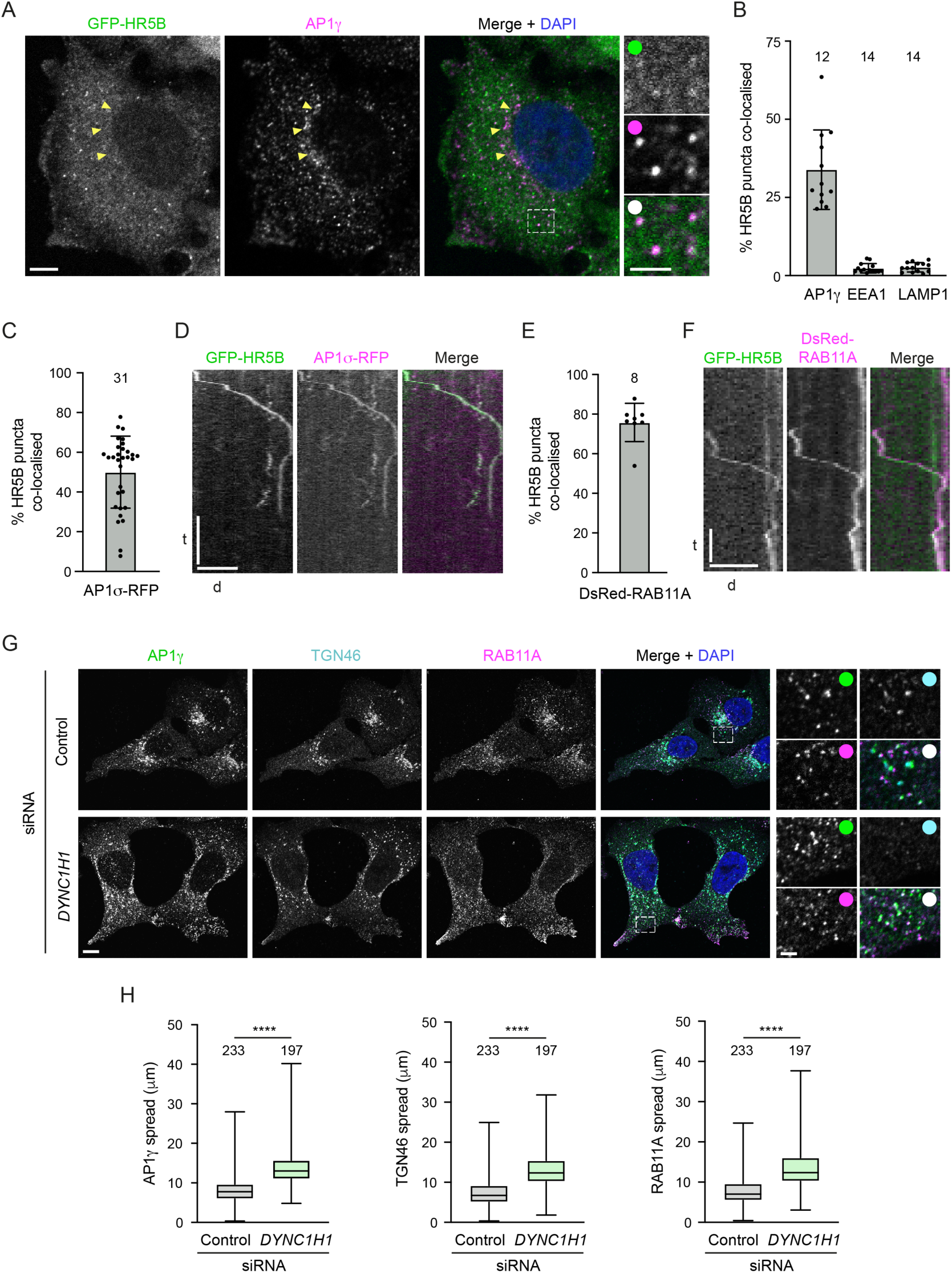
HEATR5B is co-transported with AP1 and RAB11A. **(A)** Representative confocal images of immunostained HeLa cells showing localisation of GFP-HEATR5B (HR5B) and AP1γ (GFP signal was amplified with GFP antibodies). Yellow arrows, position of TGN-associated AP1γ (which rarely co-localises with GFP-HR5B); dashed box shows area containing multiple co-localisation events that is magnified in right-hand images. In this and other figures, DAPI is used to stain DNA. **(B)** Quantification of percentage of GFP-HR5B puncta co-localised with indicated proteins in fixed cells. **(C)** Quantification of percentage of GFP-HR5B puncta that co-localise with AP1σ1-RFP in live HeLa cells. **(D)** Example kymograph (time-distance plot) of a long-range co-transport event of GFP-HR5B and AP1σ1-RFP. **(E)** Quantification of percentage of GFP-HR5B puncta that co-localise with DsRed-RAB11A in live HeLa cells. **(F)** Example kymograph of a long-range co-transport event of GFP-HR5B and DsRed-RAB11A. **(G, H)** Representative confocal images (G) and quantification (H) of spread of the indicated protein signals away from the perinuclear region in immunostained U2OS cells ± *DYNC1H1* siRNA. In A and G, as well as in other figures with insets, white circles indicate merge of magnified images. In D and F, d = distance and t = time. Scale bars: A, 10 μm; A insets, 2 μm; D, 20 μm and 20 s; F, 10 μm and 10 s; G, 10 μm.; G insets 2.5 μm. In B, C, E and H, number of cells analysed is shown above columns. In B, C, and E, circles indicate values from individual cells, with columns and error bars representing mean ± S.D.. In H, boxes show interquartile range (25^th^-75^th^ percentile of values) and horizontal line is the median; statistical significance was evaluated with a t-test: ****, p <0.0001.

We then tested if the cytoplasmic structures containing HEATR5B and AP1 are motile by live imaging of GFP-HEATR5B HeLa cells transfected with a plasmid coding for an RFP-tagged σ1 subunit of AP1. As expected from our immunostaining results, HEATR5B and AP1σ1 were frequently enriched together on punctate structures in the cytoplasm (Figure 3C, Figure S3A and Movie S1). During several minutes of filming, the vast majority of the dual-labelled punctate structures exhibited short, oscillatory movements (Movie S1). This behaviour is reminiscent of the behaviour of early endosomal cargoes for microtubule motors in HeLa cells, which can take tens of minutes to traverse the cytoplasm (Flores-Rodriguez *et al*., 2011; Tirumala *et al*., 2022; Zajac *et al*., 2013). However, a small fraction of AP1σ1 puncta exhibited unidirectional movements of between 1.5 and 10 μm both towards and away from the perinuclear region with a mean instantaneous velocity of 260 ± 4 nm/s (± SEM; 22 particles) (Figure 3D and Movie S2). Thus, AP1-associated membranes can be subjected to long-range transport.

AP1 is implicated in the anterograde movement of cargoes from the TGN to recycling endosomes, as well as a reverse process that retrieves unbound receptors and SNAREs back to the TGN in order to sustain anterograde trafficking (Cancino *et al*., 2007; Hirst *et al*., 2012; Robinson *et al*., 2010). To confirm that motile HEATR5B-positive structures in the cytoplasm are associated with recycling endosomal membranes, we transfected GFP-HEATR5B HeLa cells with a DsRed-tagged version of the recycling endosome marker RAB11A. HEATR5B and RAB11A signals frequently overlapped in the cytoplasm (Figure 3E and Figure S3B). As was the case with AP1 and GFP-HEATR5B, most of the structures positive for RAB11A and GFP-HEATR5B exhibited oscillatory motion, with only a small fraction undergoing long-distance transport (Figure 3F, Movies S3 and S4; mean instantaneous velocity of 264 ± 8 nm/s (± SEM; 15 particles)).

The above observations indicate that HEATR5B can associate with AP1-bound endosomal membranes that are capable of directed movement. To determine if dynein contributes to trafficking of these structures, we examined the distribution of AP1γ in cells treated with an siRNA pool that depletes DYNC1H1 (Figure S3C). As microtubule minus ends are enriched at the perinuclear microtubule-organising centre (Brinkley, 1985), a role of dynein in AP1 transport should be reflected in more peripheral AP1γ localisation when the motor complex is inhibited. This is indeed what we saw, with AP1γ-associated structures more dispersed in the *DYNC1H1* siRNA conditions than in controls treated with a non-targeting control siRNA pool (Figure 3G, H). The dispersed AP1γ-associated structures included TGN material (as judged by strong TGN46 staining), which was previously shown to depend on dynein for perinuclear clustering (Burkhardt *et al*., 1997). However, AP1γ puncta that were positive for RAB11A but lacked robust TGN46 signals, and thus corresponded to the free recycling endosome compartment (Fujii *et al*., 2020a), were also localised more peripherally when DYNC1H1 was depleted (Figure 3G). Based on these data, we conclude that dynein promotes retrograde trafficking of AP1-associated endosomal membranes.

### HEATR5B promotes membrane localisation and motility of AP1

We next sought to determine if HEATR5B contributes to trafficking of AP1-associated membranes. To this end, we used CRISPR/Cas9-mediated mutagenesis to generate clonal human U2OS cell lines with frameshift mutations in the *HEATR5B* gene that disrupt protein expression (Figure S4A, B, and Table S3).

We first used immunostaining to examine the effect of disrupting HEATR5B on the distribution of AP1 in fixed cells. Compared to parental, wild-type cells, *HEATR5B* mutant cells had a striking reduction in the number of AP1γ puncta in the cytoplasm, as well as lower intensity of the puncta that were present (Figure 4A – C). This phenotype was seen in three independent *HEATR5B* mutant U2OS clones and was fully rescued by transfection of a GFP-HEATR5B construct (Figure S5 and S6A), confirming the causal nature of the *HEATR5B* mutation. To better understand the nature of the phenotype, we visualised RAB11A together with AP1γ in control and mutant cells (Figure 4A). As expected, in the control cells punctate signals of both proteins frequently overlapped with each other within the cytoplasm. RAB11A and AP1γ signals did not, however, co-localise precisely, in keeping with a report that AP1 is present on tubular endosomes that have different constituents spatially segregated (Klumperman & Raposo, 2014). In *HEATR5B* mutant cells, RAB11A-positive structures were still abundant. However, there was a strong reduction in the amount of AP1γ signal associated with them.

**Figure 4.**
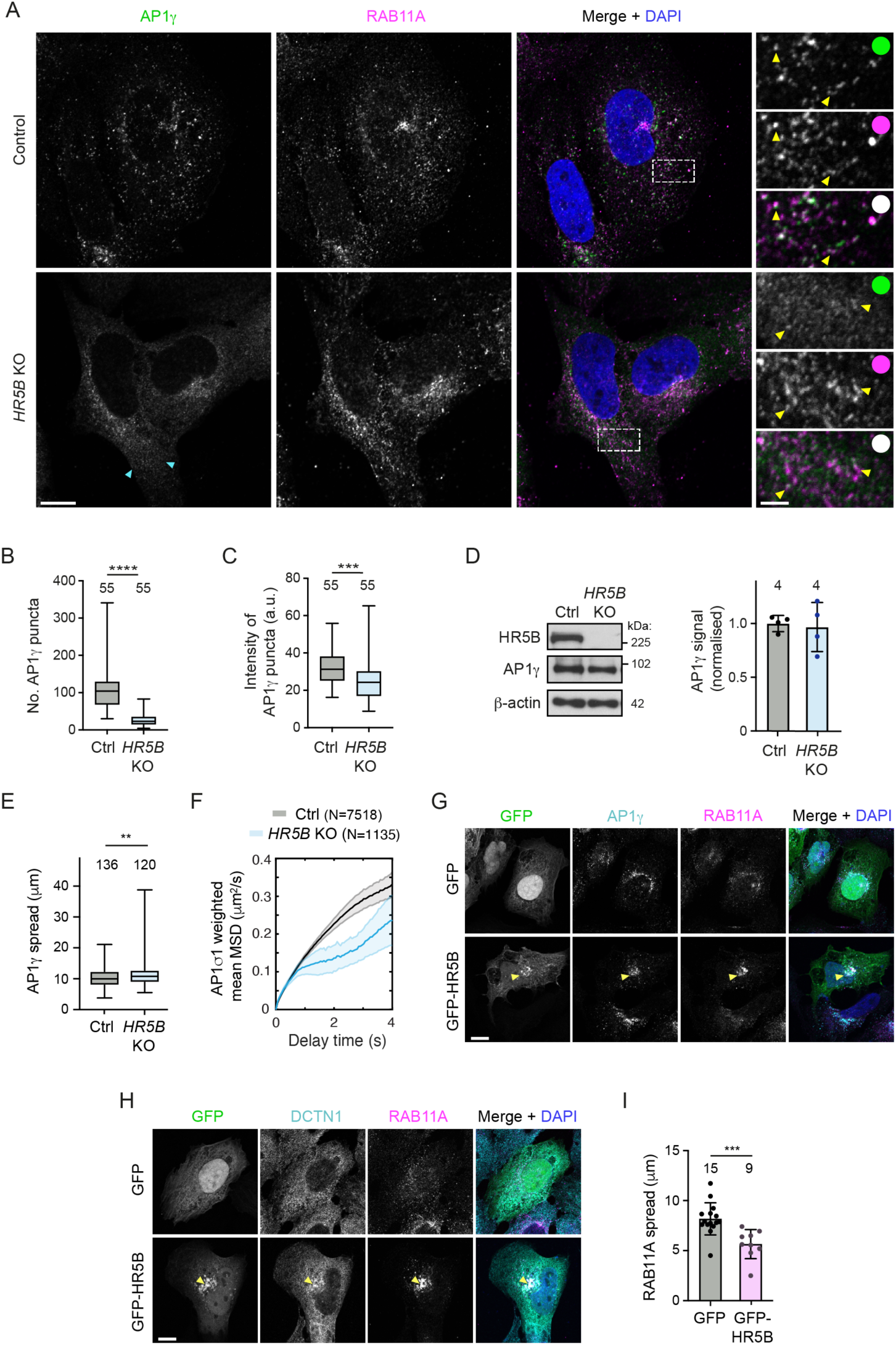
HEATR5B promotes AP1 membrane localisation and motility. **(A)** Representative confocal images of AP1γ and RAB11A in immunostained wild-type (control) and *HEATR5B* mutant (*HR5B* KO) U2OS cells. Dashed box shows region magnified in right-hand images. Blue arrowheads show diffuse AP1γ signal in the cytoplasm, which is quantified in addition to punctate signal. Yellow arrowheads in insets show examples of AP1γ association with RAB11A. **(B, C)** Quantification of number (B) and mean total intensity (C) of AP1γ puncta in control (Ctrl) and *HR5B* KO U2OS cells (a.u., arbitrary units). **(D)** Left, immunoblot images showing levels of HR5B and AP1γ in control and *HR5B* KO U2OS cells (β-actin, loading control). Right, quantification of AP1γ signal (normalised to β-actin signal). **(E)** Quantification of spread of total AP1γ signal away from the perinuclear region in control and *HR5B* KO U2OS cells. **(F)** Quantification of weighted mean MSD of AP1σ1-RFP puncta in control and *HR5B* KO U2OS cells. **(G, H)** Representative confocal images showing localisation of RAB11A (G and H), AP1γ (G) and DCTN1 (H) in cells that strongly express GFP-HR5B or a GFP control. Arrowheads show perinuclear clustering of signals in GFP-HR5B cells. **(I)** Quantification of spread of RAB11A signal away from perinuclear region in GFP or GFP-HR5B overexpressing U2OS cells. Scale bars: A, 10 μm; A insets 2.5 μm; G and H, 5 μm. In B, C, and E, boxes show interquartile range (25^th^-75^th^ percentile of values) and horizontal line is the median. In I, circles indicate values from individual cells; columns and error bars represent mean ± S.D.. Number of cells (in B, C, E and I) or lysates (D) analysed is shown above columns. In F, N = number of particles (from 31 control and 17 KO U2OS cells). Statistical significance was evaluated with a t-test: ****, p <0.0001; ***, p <0.001; **, p <0.01.

These data indicate that disrupting HEATR5B reduces the association of AP1γ with endosomal membranes. We also observed modestly reduced association of AP1γ with the TGN in the *HEATR5B* mutant cells, which was rescued by transfection of the GFP-HEATR5B construct (Figure S6A, B). Consistent with reduced AP1 interaction with the recycling compartment, disruption of HEATR5B caused excessive tubulation of Transferrin receptor-positive membranes that were associated with the TGN (Figure S7A, B). The decreased interaction of AP1γ with endosomal membranes and the TGN in mutant U2OS cells is consistent with the reduction in punctate AP1 signal observed when the HEATR5B orthologue was knocked down with RNAi in *C. elegans* (Gillard *et al*., 2015) and *Drosophila* wing discs (Le Bras *et al*., 2012). However, these earlier studies did not determine if targeting HEATR5B affects expression, stability or membrane recruitment of AP1. Immunoblotting of extracts showed that the overall level of AP1γ protein was not altered in the *HEATR5B* mutant U2OS cells (Figure 4D). We therefore conclude that HEATR5B promotes recruitment of AP1 to endosomal membranes and, to a lesser extent, the TGN.

Our previous observation that bright AP1γ puncta are abundant in cells treated with *DYNC1H1* siRNA (Figure 3G) revealed that HEATR5B does not co-operate with dynein to promote AP1 membrane localisation. However, a small but significant dispersion of total AP1γ signal towards the periphery of *HEATR5B* mutant U2OS cells (Figure 4E) was consistent with HEATR5B having an additional function in dynein-based motility of AP1-bound membranes. To directly assess the contribution of *HEATR5B* to motility of AP1, we performed high-speed imaging of AP1σ1-RFP in live wild-type and mutant U2OS cells (Movie S5). As expected from our fixed cell analysis, AP1σ1 puncta were dimmer in the HEATR5B deficient cells than in controls (Figure S7C). In both genotypes, the low intensity of AP1σ1-RFP puncta meant that many of these structures could only be followed for a few seconds before the signals bleached. Nonetheless, mean square displacement (MSD) analysis over this timescale revealed that the motility of AP1σ1 puncta in the cytoplasm of mutant cells was much less persistent than in controls (Figure 4F). The impaired transport of AP1-positive structures when *HEATR5B* was disrupted was not an indirect effect of reduced AP1 association with membranes, as the motility defect was still evident for AP1σ1 puncta that had equivalent intensities in control and *HEATR5B* mutant cells (Figure S7C). Staining of *HEATR5B* deficient cells with an antibody to α-Tubulin indicated that altered AP1σ1 motility was also not due to impaired integrity of the microtubule network (Figure S7D). Collectively, these data indicate that HEATR5B directly promotes motility of AP1-positive structures in U2OS cells.

We next asked if HEATR5B is sufficient to redistribute AP1-associated membranes by strongly expressing GFP-HEATR5B in U2OS cells via transfection. Compared to control cells in which only GFP was expressed, GFP-HEATR5B expressing cells had increased perinuclear clustering of RAB11A-associated membranes that were also positive for AP1γ and the DCTN1 subunit of the dynactin complex (Figure 4G – I). These observations suggest that HEATR5B can stimulate retrograde trafficking of AP1-associated endosomal membranes by dynein-dynactin.

### The *Drosophila HEATR5B* homologue is an essential gene

Our experiments in human tissue culture cells revealed that HEATR5B promotes recruitment of AP1 to endosomal membranes, as well as the motility of these structures. To assess the importance of HEATR5B function at the organismal level, as well as in polarised cell types, we generated a strain of the fruit fly *Drosophila melanogaster* with an early frameshift mutation in the single *HEATR5B* homologue (*CG2747*, hereafter called *Heatr5*) (Figure S8A). This was achieved by combining a Cas9 transgene that is active in the female germline (*nos-cas9*) with a transgene expressing two gRNAs that target *Heatr5* (*gRNA-Hr5^1+2^*) (Figure S8A). Zygotic *Heatr5* homozygous mutants (*Hr5^1^/Hr5^1^*) failed to reach adulthood (Figure 5A), with most animals dying during the 2nd larval instar stage (Figure S8B). The lethal phenotype was not complemented by a pre-existing deletion of a genomic region that includes the *Heatr5* gene (Figure S8B) but was fully rescued by a wild-type Heatr5 transgene (Figure 5A). Thus, the lethality observed was due to the *Heatr5* mutation and not an off-target effect of the gRNAs.

**Figure 5.**
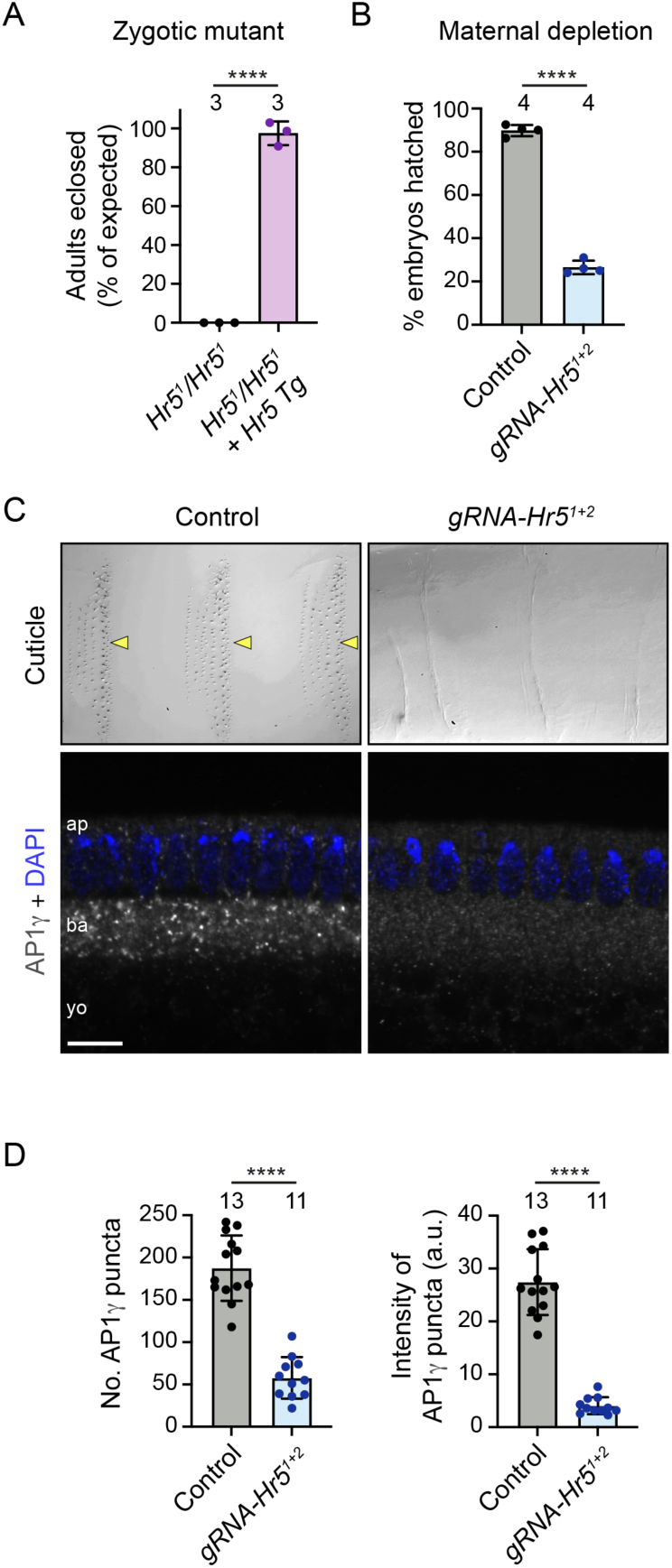
*Drosophila* Heatr5 is required for viability and promotes membrane association of AP1. **(A)** Quantification of survival to adulthood of zygotic *Heatr5* (*Hr5*) homozygous mutants in the absence and presence of a wild-type *Hr5* transgene (Tg). **(B)** Quantification of hatching of eggs laid by control (*nos-cas9*) and *nos-cas9 gRNA-Hr5^1+2^* females. **(C)** Representative images of cuticle preparations of late-stage embryos (top panels) and AP1γ distribution in blastoderm-stage embryos (bottom panels) laid by control and *nos-cas9 gRNA-Hr5^1+2^* females. In C, arrowheads point to denticle belts in the control, and ‘ap’, ‘ba’ and ‘yo’ refer to apical cytoplasm, basal cytoplasm and yolk, respectively. Scale bar: top panels, 30 μm; bottom panels 10 μm. **(D)** Quantification of number and mean total intensity of AP1γ puncta in blastoderm embryos laid by control and *nos-cas9 gRNA-Hr5^1+2^* mothers. In A, B and D, circles indicate values from individual crosses (A, B) or embryos (D); columns and error bars represent mean ± S.D.; number of independent crosses or embryos analysed is shown above columns (in A and B, at least 200 flies or embryos were analysed per cross); statistical significance was evaluated with a t-test: ****, p <0.0001.

To examine the maternal requirement for Heatr5, we followed the development of the embryos laid by *nos-cas9 gRNA-Hr5^1+2^* mothers. The vast majority of embryos did not hatch into larvae (Figure 5B), instead arresting during late embryogenesis. These embryos, which presumably had biallelic disruption of *Heatr5* in the female germline, typically had denticle hairs that were either absent or short and thin (Figure 5C and Figure S8C). This phenotype is reminiscent of that of mutants for Syntaxin-1A, which promotes apical protein secretion (Moussian *et al*., 2007; Schulze & Bellen, 1996). Taken together, these results demonstrate that Heatr5 has essential zygotic and maternal functions in *Drosophila*.

### Heatr5 strongly promotes dynein-based transport of AP1-positive structures in the fly embryo

We next set out to understand the effect of disrupting Heatr5 on AP1-based trafficking in *Drosophila*. For these experiments, we used the syncytial blastoderm embryo as a model. This system is attractive because the microtubule cytoskeleton is highly polarised with minus ends nucleated apically above the nuclei and plus ends extended basally (Karr & Alberts, 1986; Warn & Warn, 1986). This means that the activity of dynein and kinesin motors can be distinguished by the direction of cargo movement (Shubeita *et al*., 2008). Moreover, membranes can be readily visualised by microinjection of antibodies coupled to bright fluorophores into the shared cytoplasm of the syncytium (Papoulas *et al*., 2005; Sisson *et al*., 2000).

We first analysed the effect of Heatr5 depletion on AP1 distribution in blastoderm embryos by immunostaining embryos laid by control and *nos-cas9 gRNA-Hr5^1+2^* mothers with antibodies to AP1γ. Bright AP1γ puncta were abundant in the cytoplasm of control embryos, particularly in the region basal to the nuclei (Figure 5C). In contrast, embryos of *nos-cas9 gRNA-Hr5^1+2^* females had AP1γ puncta that were much fewer in number and much dimmer (Figure 5C, D), reminiscent of the situation in *HEATR5B* deficient human cells. This phenotype was confirmed with an independent pair of *Heatr5* gRNAs (Figure S9A, B), as was the failure of mutant embryos to develop to larval stages (Figure S9C). As in *HEATR5B* deficient human cells, the change in AP1γ distribution in *nos-cas9 gRNA-Hr5* embryos was not due to altered total amounts of AP1γ protein (Figure S9D). These observations reveal a conserved role of HEATR5B proteins in localising AP1 to membranes.

We next examined motility of AP1-positive structures in live embryos. This was achieved by labelling the AP1γ antibodies with Alexa555-coupled secondary antibodies, injecting the conjugates into the embryo at the junction of the yolk and basal cytoplasm (Figure S10A), and filming the peripheral region of the embryo for several minutes. Following injection of antibody conjugates into control embryos, fluorescent puncta formed that underwent rapid, bidirectional transport in the cytoplasm. These movements had a strong net apical bias, often pausing or arresting just beneath the peripheral blastoderm nuclei (Figure 6A, Figure S10B and Movie S6). Punctate signals were not observed with fluorescent secondary antibodies injected alone or when bound to control primary antibodies (Figure S10C), confirming their dependence on the AP1γ antibody.

**Figure 6.**
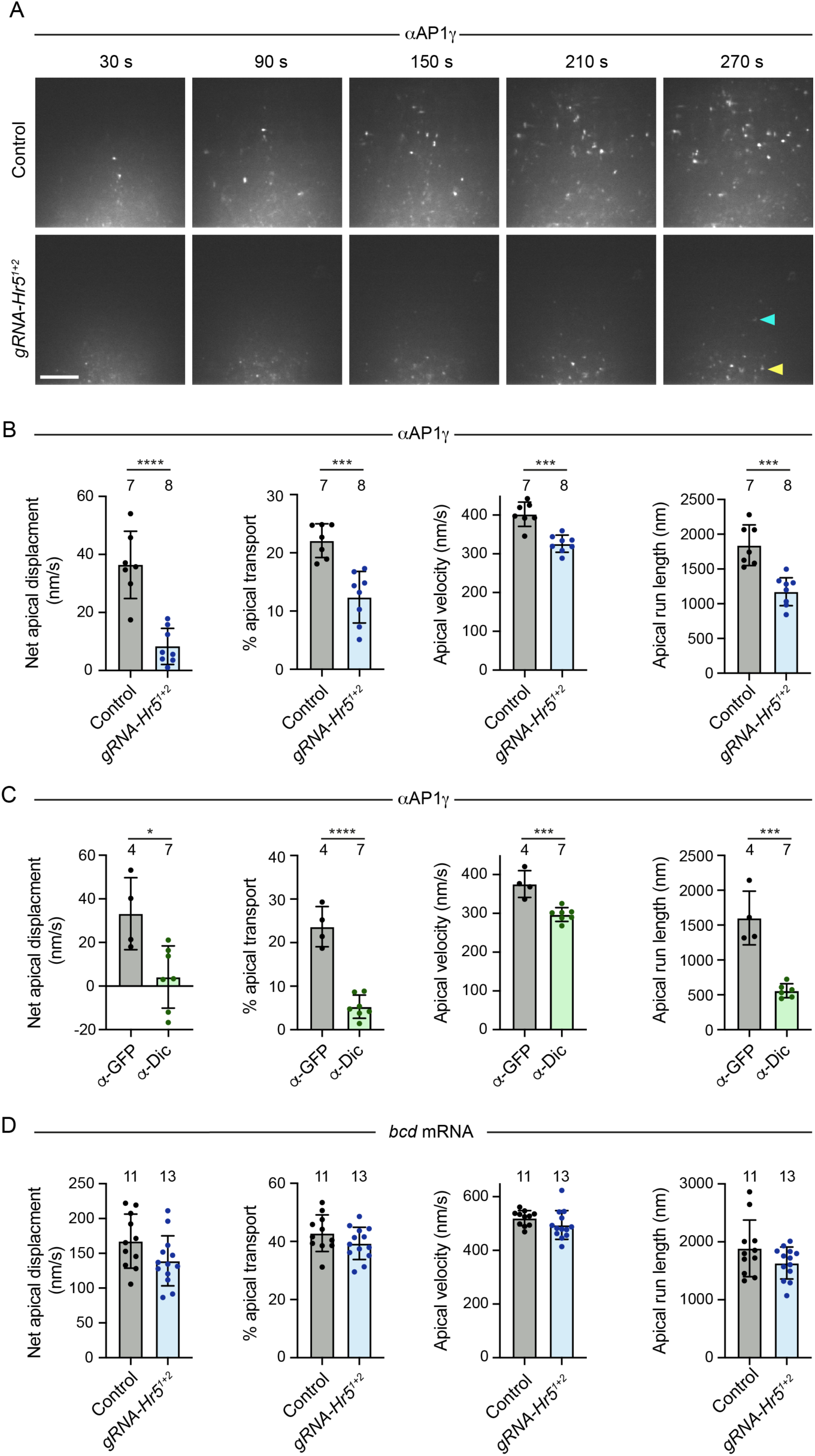
Heatr5 promotes dynein-based motility of AP1-positive structures in the blastoderm embryo. **(A)** Stills from representative image series of blastoderm embryos from control (*nos-cas9*) and *nos-cas9 gRNA-Hr5^1+2^* mothers injected at the junction of the yolk and basal cytoplasm with fluorescent AP1γ antibody conjugates. Yellow arrowhead, basally concentrated puncta in mutant embryo; blue arrowhead, rare punctum that underwent long-range apical transport in mutant embryo (see Movie S6). Scale bar, 10 μm. **(B – D)** Quantification of motility of AP1γ (B, C) and injected *bcd* mRNA (D) in embryos of control and *nos-cas9 gRNA-Hr5^1+2^* mothers (B, D) or wild-type embryos pre-injected with function-blocking Dic antibodies or control GFP antibodies (C). ‘% apical’ is the percentage of the particle trajectory time that is classed as apical transport. Circles are mean values for individual embryos; columns and error bars represent means ± S.D. of these mean values; number of embryos analysed is shown above columns (at least 24 particles analysed per embryo); statistical significance was evaluated with a t-test: ****, p < 0.0001; ***, p <0.001; *, p <0.05.

We next injected the fluorescent AP1γ antibody conjugates into *nos-cas9 gRNA-Hr5^1+2^*embryos. As expected, the AP1γ puncta in these embryos were considerably dimmer than those observed in the control. Nonetheless, the antibody labelling method meant they were bright enough to be followed throughout the period of filming. In the mutant embryos, the rate of net apical movement of AP1γ puncta was strongly impaired. The puncta mostly exhibited short-range saltatory movements or pausing behaviour (Figure S10B and Movie S6) and consequently rarely reaching the region beneath the blastoderm nuclei during the period of image acquisition (Figure 6A). A defect in the rate of net apical transport in mutant embryos was confirmed by automated tracking of particle movement, which additionally revealed significant reductions in the apical velocity and run length of AP1γ signal compared to the control (Figure 6B), as well as more modest decreases in the basal velocity and run length (Figure S11A). These observations show that AP1 undergoes net apical transport in the *Drosophila* embryo and that this process is strongly promoted by Heatr5.

As actin structures are concentrated above the nuclei at blastoderm stages (Karr & Alberts, 1986), it is likely that microtubules are the tracks for long-distance transport of AP1 in the basal cytoplasm. Supporting this notion, AP1γ transport in wild-type embryos was arrested by microinjection of the microtubule targeting agent colcemid and rapidly reinitiated when the drug was inactivated with a pulse of UV light (Movie S7) (Czaban & Forer, 1985). The localisation of microtubule minus ends above the blastoderm nuclei strongly suggests that apical AP1 transport is driven by dynein and we confirmed this is the case by injecting wild-type embryos with a function-blocking antibody to Dynein intermediate chain (Dic) (Bullock *et al*., 2006) prior to injecting the AP1γ antibody conjugate (Figure 6C and Movie S8). Consistent with the interdependence of dynein and kinesin motors in several bidirectional transport systems (Jolly & Gelfand, 2011), the Dic antibody also impaired some features of plus-end-directed AP1γ motility (Figure S11B). Thus, the modest impairment of plus-end-directed motion of AP1γ in *Heatr5* mutant embryos could be an indirect effect of dynein inhibition. Together, these experiments demonstrate that Heatr5 promotes dynein-dependent transport of AP1 along microtubules.

To test if Heatr5 has a general role in dynein-based transport in the embryo, we assessed mRNA trafficking by the motor in wild-type and *nos-cas9 gRNA-Hr5^1+2^* embryos via microinjection of fluorescent *bicoid* (*bcd*) RNA (Bullock & Ish-Horowicz, 2001; Bullock *et al*., 2006; Snee *et al*., 2005; Wilkie & Davis, 2001). Neither apical nor basal mRNA transport was significantly impaired by disruption of Heatr5 (Figure 6D, Figure S11C and Movie S9). Thus, Heatr5 selectively promotes dynein-mediated transport of AP1-positive structures in the embryo.

### Heatr5-dependent AP1 trafficking routes in the embryo involve endosomal and Golgi membranes

To next sought to shed light on the trafficking routes of the transported AP1-associated structures in the embryo. To this end, we used CRISPR-mediated homology-directed repair to generate a fly strain in which the *trans*-Golgi marker Golgin-245 is endogenously tagged with GFP (Figure S12) and confirmed that the fusion protein is correctly localised to the dispersed ‘mini-stacks’ that constitute the Golgi apparatus in *Drosophila* cells (Kondylis & Rabouille, 2009) (Figure S13A). Injecting the fluorescent AP1γ antibody conjugate into the GFP-Golgin-245 embryos revealed that AP1 puncta were often transported to, and engaged with, the periphery of Golgin-245-positive structures or were trafficked together with them (Figure 7A, Figure S13B and Movie S10 – 12). Consistent with these observations, AP1γ puncta were frequently located adjacent to Golgin-245 puncta in fixed, uninjected embryos (Figure S13C). These findings suggest that microtubule-based transport of AP1-associated membranes facilitates their interaction with Golgi membranes.

**Figure 7.**
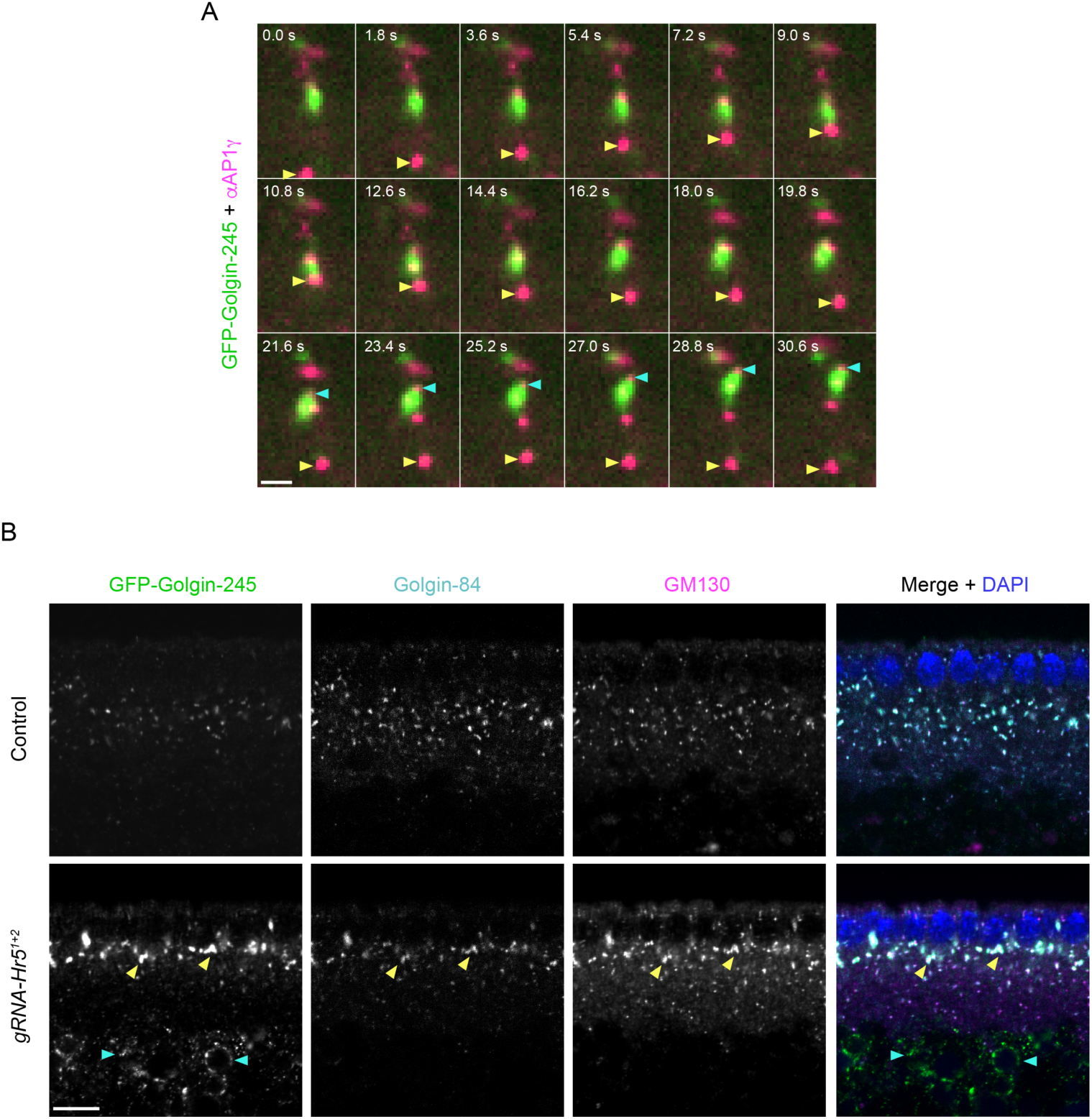
*Drosophila* Heatr5 promotes transport of AP1-positive structures to the Golgi and is required for Golgi organisation. **(A)** Stills from representative image series of GFP-Golgin-245 knock-in embryos injected with fluorescent AP1γ antibody conjugates. Yellow arrowhead shows an AP1γ-positive structure that is transported apically, interacts with a Golgi body and is then transported basally. Blue arrowhead shows AP1γ at the front of an apically transported Golgi body. **(B)** Representative confocal images of GFP-Golgin-245, Golgin-84 and GM130 distribution in embryos from control (*nos-cas9*) and *nos-cas9 gRNA-Hr5^1+2^* mothers. Yellow and blue arrowheads show, respectively, examples in mutant embryo of expanded Golgi structures beneath the nuclei and ectopic accumulation of GFP-Golgin-245 in the yolk. Scale bars: A, 2 μm; B, 10 μm.

Staining of fixed YFP-Rab11 knock-in embryos (Dunst *et al*., 2015) demonstrated that many of the AP1γ-positive structures in the basal cytoplasm associated with Rab11 (Figure S13D). However, AP1γ and Rab11 signals rarely overlapped precisely. This finding suggests that, as in U2OS cells (Figure 4A), these proteins are present on closely opposed or conjoined endosomal membrane structures. In contrast, AP1γ was not enriched in the vicinity of the pool of Rab11 that is located at the apically positioned microtubule-organising centre (Pelissier *et al*., 2003) (Figure S13D). These observations implicate the transported AP1 structures in trafficking events in the basal cytoplasm involving membranes of the recycling endosome compartment and the Golgi apparatus.

### Heatr5 is required for Golgi organisation in the *Drosophila* embryo

Finally, we investigated if the AP1 membrane recruitment and trafficking defects in Heatr5-deficient embryos are accompanied by defects in Golgi morphology. We compared the distribution in control and *Heatr5* mutant embryos of the Golgin-245, GM130 and Golgin-84 golgin proteins, which are enriched, respectively, in the *trans*-Golgi, *cis*-Golgi and rims of the Golgi stack (Kondylis & Rabouille, 2009; Munro, 2011). In contrast to the small Golgi stacks distributed throughout the basal cytoplasm in the wild-type embryo, the *Heatr5* mutant embryos had large conglomerations of all three proteins in the region of the basal cytoplasm just beneath the nuclei (Figure 7B; quantification in Figure S14). Of the three Golgi proteins analysed, Golgin-245 showed a particularly strong conglomeration phenotype in the mutant.

Increased aggregation of Golgi stacks has previously been observed in *Drosophila* S2 cells treated with brefeldin A (Fujii *et al*., 2020b), which impairs AP1 association with membranes by targeting ARF1 function (Donaldson *et al*., 1992; Helms & Rothman, 1992). Conglomeration of golgin signals in *Heatr5* mutant embryos may therefore reflect impaired release of material from the Golgi due to inefficient targeting of AP1 complexes to this organelle. Based on these observations, we propose that Heatr5-mediated transport of AP1 complexes to the Golgi stimulates post-Golgi trafficking in the embryo. We additionally discovered that a fraction of Golgin-245 was ectopically localised in the yolk in *Heatr5* mutant embryos (Figure 7B; quantification in Figure S14). This result raises the possibility that Heatr5 also contributes to transport of this protein from an internal pool to the cytoplasm.

## DISCUSSION

### Novel interactors of the dynein tail

In addition to known co-factors, our recombinant tail-based screening strategy revealed ∼50 novel candidate interactors of the dynein complex in brain extracts. Several of these proteins had their association with the tail enhanced by dynactin. The novel dynein tail interactors may not have been found in previous dynein ‘interactomes’ (Gershoni-Emek *et al*., 2016; Redwine *et al*., 2017) because these studies used full-length DYNC1H1, in which the presence of the motor domain can impair cargo association with the tail (Htet *et al*., 2020; Zhang *et al*., 2017), as well as different cell types or subcellular compartments as a source of potential interactors. Sequence analysis revealed that only ∼20% of the total set of dynein-interacting proteins in our experiments contain a predicted coiled-coil domain (Tables S1 and S2), suggesting that many of them interact indirectly with the motor complex or bind directly through a mode distinct from that of canonical activating adaptors. Whilst we prioritised HEATR5B – one of the dynactin-stimulated interactors – for mechanistic analysis in this study, we anticipate that investigating other hits from our screen will shed further light on dynein’s cargo linkage and regulation. Of particular note, we observed a large number of ribosomal proteins and other RNA-associated proteins in our dynein tail interactomes. These factors are candidates to participate in dynein-mediated trafficking of messenger ribonucleoprotein particles (mRNPs), a process that can dictate the site of protein function in cells (Mofatteh & Bullock, 2017). Other intriguing hits include the dynactin-stimulated tail interactor Wdr91, a Rab7 effector implicated in endosomal recycling and lysosomal function (Liu *et al*., 2022; Xing *et al*., 2021), as well as reovirus infection (Snyder *et al*., 2022). In addition to analysing the current set of dynein interactors, it may be possible in the future to adapt our pulldown approach to identify more transient interactors of the motor complex, for example by incorporating a biotin ligase on the dynein tail for proximity-dependent biotinylation (Samavarchi-Tehrani *et al*., 2020).

### HEATR5B promotes AP1 membrane localisation and motility

HEATR5B was first identified in human cells through its association with AFTPH, which contains a motif that binds the AP1γ ‘ear’ domain (Hirst *et al*., 2005; Lui *et al*., 2003). It was shown that AFTPH and HEATR5B form a stable complex with SYNRG and that knocking down the function of this assembly causes AP1 cargoes to be partially re-routed from the TGN to a more peripheral compartment (Hirst *et al*., 2005). RNAi-based knockdowns subsequently implicated HEATR5B orthologues in AP1-based trafficking of components of the Notch signalling pathway in *Drosophila* imaginal discs (Le Bras *et al*., 2012) and E-cadherin in *C. elegans* epidermal cells (Gillard *et al*., 2015). How HEATR5B mechanistically influences AP1 function in these systems was not, however, investigated.

It was previously shown that the budding yeast homologue of Heatr5B – Laa1 – is required for membrane recruitment of AP1 (Fernandez & Payne, 2006; Zysnarski *et al*., 2019). We have shown that HEATR5B proteins also promote association of AP1 with membranes in human and *Drosophila* cells. Thus, HEATR5B has widespread roles in membrane targeting of AP1. In budding yeast, the ability of Laa1 to promote AP1 membrane association depends on another protein, Laa2 (Zysnarski et al., 2019). It is unclear how HEATR5B promotes AP1 membrane association in higher eukaryotes, as an overt orthologue of Laa2 is missing. However, our data indicate that this process is independent of HEATR5B’s ability to bind dynein. We provide evidence that HEATR5B’s interaction with the motor complex instead promotes microtubule-based trafficking of AP1-associated endosomal membranes. Thus, HEATR5B has a central role in controlling the function of AP1, co-ordinating its association with membranes and microtubule-based transport of these structures.

Work by several groups has demonstrated the importance of coiled-coil-containing activating adaptors in linking dynein to cargoes and dynactin, and thus initiating long-distance cargo transport (Olenick & Holzbaur, 2019; Reck-Peterson *et al*., 2018). Whilst our data demonstrate a role for HEATR5B proteins in promoting AP1 transport, including long-range dynein-based movements in the fly embryo, several observations suggest they are unlikely to act analogously to an activating adaptor as a primary link between AP1-associated membranes and dynein. Firstly, HEATR5B proteins lack the coiled-coil domains that are typical of activating adaptors. Secondly, our observation of residual dynein-based motility of AP1-positive membranes in *nos-cas9 gRNA-Hr5 Drosophila* embryos shows the cargo can still be linked to the motor complex when HEATR5B is disrupted. And thirdly, in ongoing work, we have failed to detect *in vitro* activation of dynein-dynactin motility by purified HEATR5B. We therefore favour a scenario in which HEATR5B enhances the function of a dynein-dynactin-activating adaptor complex on AP1-associated membranes. The involvement of an as-of-yet unidentified activating adaptor may also explain why we observed stimulation of dynein’s association with HEATR5B by dynactin in cellular extracts but not with purified proteins.

HEATR5B consists almost entirely of repeats of HEAT domains, ∼40 amino acid motifs of anti-parallel α-helices separated by a short linker (Andrade & Bork, 1995; Groves *et al*., 1999). The ability of HEAT repeat proteins to act as flexible scaffolds for protein-protein interactions (Grinthal *et al*., 2010; Yoshimura & Hirano, 2016) raises the possibility that HEATR5B promotes transport by stabilising a dynein-dynactin-activating adaptor assembly and/or its association with the membrane. Intriguingly, another non-coiled-coil protein involved in cargo transport by dynein-dynactin, Ankyrin-B (Lorenzo *et al*., 2014), also contains a large number of α-helical ankyrin repeats. It is tempting to speculate that the repeated units in Ankyrin-B and HEATR5B play an analagous role in scaffolding dynein-dynactin-activating adaptor-cargo complexes.

Future efforts will be directed at identifying additional proteins that link AP1 to dynein and dynactin and determining how HEATR5B affects the motility of the entire machinery when it is reconstituted *in vitro*. It will also be important to determine if and how HEATR5B puncta contributes to transport of other cargoes by dynein. Whilst our microinjection of mRNAs in mutant fly embryos show that HEATR5B is not a general regulator of dynein activity, the finding that a substantial fraction of HEATR5B puncta in human cells do not overlap with AP1 raises the possibility of involvement in transport of additional cargoes.

### The role of HEATR5B and dynein in AP1-based membrane trafficking

Dynein plays a role in multiple trafficking events in the endocytic pathway, including transport of peripheral early endosomes, late endosomes and lysosomes, as well as sorting of internalised receptors through these compartments (Burkhardt *et al*., 1997; Driskell *et al*., 2007; Guo *et al*., 2016; Hong *et al*., 2009; Horgan *et al*., 2010; Jongsma *et al*., 2023; Jordens *et al*., 2001; Lalli *et al*., 2003; Loubery *et al*., 2008; Traer *et al*., 2007). The range of dynein functions makes it challenging to study specific trafficking processes by targeting the motor complex. Our analysis of HEATR5B highlights a novel dynein-based process for retrograde trafficking of AP1-associated endosomal material to the Golgi apparatus (Hirst *et al*., 2012; Robinson *et al*., 2010). This process appears to be distinct from a previously identified dynein-and RAB11FIP3-dependent process for moving material between RAB11A-positive recycling endosomes and the the TGN (Horgan *et al*., 2010; McKenney *et al*., 2014) because RAB11A and AP1 are not enriched on the same domain of tubular endosomal structures. We envisage the two dynein-based pathways acting in parallel to ensure efficient delivery of endosomal material to the TGN or translocating distinct sets of proteins that are sorted into endosomal membrane domains enriched with AP1 or RAB11A. Our observation in human cells that HEATR5B is concentrated on peripheral AP1-positive endosomal membranes but not the perinuclear AP1-positive TGN additionally suggests a mechanism for locally modulating dynein-based transport. Limiting dynein activity on AP1-bound membranes at the TGN presumably facilitates kinesin-driven trafficking of AP1 cargoes in the anterograde direction (Nakagawa *et al*., 2000; Schmidt *et al*., 2009), thus ensuring bidirectional trafficking.

In addition to trafficking cargoes, the HEATR5B-mediated transport process may promote delivery of the AP1 complex from endosomal membranes to the TGN, where it is needed for clathrin-mediated budding of post-Golgi membranes. In support of this notion, we observed long-range transport of AP1-associated membranes from the basal cytoplasm to Golgi stacks in the wild-type *Drosophila* embryo. Moreover, impairment of this process in *Heatr5* mutant embryos was accompanied by reduced Golgi association of AP1 and a large increase in the size of the *trans*-Golgi compartment. Our observations of partially reduced AP1 localisation with the TGN and excessive tubulation of TGN-associated membranes in HEATR5B deficient human cells is consistent with dynein-mediated delivery of AP1 complexes contributing to post-Golgi trafficking in other systems. To what extent defects in this and other aspects of AP1 trafficking are involved in the neurodevelopmental syndrome associated with hypomorphic *HEATR5B* mutations (Ghosh *et al*., 2021) is another important question to address in the future.

## MATERIAL AND METHODS

### Cell culture

*Spodoptera frugiperda* (fall armyworm) Sf9 insect cells (Oxford Expression Technologies Ltd) were cultured at 27°C in Insect-XPRESS protein-free insect medium with L-Glutamine (Lonza) in shaking suspension. HEK293 Flp-in cells (provided by A. Castello, MRC-University of Glasgow Centre for Virus Research, UK), HeLa Flp-In cells ((Kaiser *et al*., 2008); provided by E. Dobrikova and M. Gromeier, Duke University Medical Center, USA), and unmodified U2OS cells and HeLa cells (both provided by H. McMahon, MRC-LMB, Cambridge, UK) were cultured at 37°C with 5% CO_2_ in complete DMEM (high glucose DMEM + GlutaMax (Gibco), 10% foetal bovine serum (Gibco) and 1% Penicillin/Streptomycin solution (Gibco)). Flp-In cells were maintained in the presence of 100 μg/ml zeocin and 5 μg/ml blasticidin (both from Gibco). Cells were checked for the absence of *Mycoplasma* using the MycoAlert kit (Lonza) at the onset of the study.

### Plasmids

pIDC-LIC2-IC2C-Tctex1-Robl1-LC8 (Schlager *et al*., 2014) and pACEBac1-His-ZZ-LTLTL-DYNC1H1E1074-GST (for generating baculovirus for expression of the dynein complex in Sf9 cells) were provided by A. Carter (MRC-LMB, Cambridge, UK). pcDNA3.1-eGFP-HR5B and pcDNA5-FRT/TO-eGFP-HR5B (for, respectively, transient or tetracyclin-inducible expression of human HEATR5B with an N-terminal GFP tag in human cells) were cloned by Gibson assembly with the full-length human HEATR5B open reading frame (derived from plasmid RC22610 (Origene)) and either pcDNA3.1-eGFP-linker or pcDNA5-FRT/TO-eGFP-linker plasmids (coding for eGFP and a GGSGGSGG linker; provided by A. Castello). pOG44 (Invitrogen) was used for expression of Flp recombinase in the Flp-in system. pAP1σ1-RFP (for expression of fluorescently-tagged APσ1 in human cells) was cloned by Gibson assembly using sequences derived from pAP1α1-eGFP (Addgene plasmid 53611) and pTagRFP-RAB2A (provided by S. Munro, MRC-LMB, Cambridge, UK). pDsRed-RAB11A WT (for expression of a fluorescently-tagged, wild-type version of RAB11A in human cells) was obtained from Addgene (plasmid 12679). For expression of GFP-HEATR5B from baculovirus, pACEBac1 eGFP-HR5B-PreSci-2x Strep tagII (tagged at the N-terminus with eGFP+linker and including a protease PreScission site and 2 Strep Tag II sequences at the C-terminus) was constructed by Gibson assembly using plasmids pACEBac1-G (containing the PreSci-2xStrep tagII sequence; provided by Eeson Rajendra (MRC-LMB, Cambridge, UK)), and pcDNA3.1 eGFP-HR5B. The sequences of all plasmids were confirmed by Sanger sequencing before use.

### Protein expression and purification

#### Human dynein tail complex

Baculovirus encoding the human dynein tail complex was produced using published procedures (Schlager *et al*., 2014). Briefly, pACEBac1-His-ZZ-LTLTL-DYNC1H1E1074-GST and pIDC-LIC2-IC2C-Tctex1-Robl1-LC8 were recombined by Cre-mediated fusion and incorporated into the baculovirus genome by transformation into EMBacY bacterial cells. Sf9 cells (500 mL cell suspension at 2 x 10^6^ cells/mL) were infected with baculovirus and cultured for a further 72 h before cell pelleting by centrifugation. Pellets were lysed by Dounce homogenisation in lysis buffer (25 mM Hepes pH7.2, 100 mM NaCl, 10 mM imidazole, 10% Glycerol, 1 mM DTT, 0.1 mM ATP, 1X COMPLETE protease inhibitor (Roche) and 0.8 mM PMSF). The lysate was centrifuged for 45 min at 70,000 rpm in a Ti70 rotor and the supernatant loaded in a 5-mL HisTrap Ni-NTA column using an AKTA Purifier system (Cytiva). The column was washed with eight column volumes of 10% (v/v) elution buffer (lysis buffer containing 500 mM imidazole) and the protein complex eluted using a step gradient of elution buffer (from 10% to 40%). The relevant fractions were collected, pooled and filtered through a 0.22-μm filter before ion exchange purification on a MonoQ column (10 mL, 5/50 GL; Sigma) that was pre-equilibrated in lysis buffer. After washing with ten column volumes of 10% buffer B (lysis buffer including 1M imidazole), the protein complex was eluted using a step gradient of buffer B (from 10% to 50%). Fractions were collected and analysed by SDS-PAGE, followed by pooling of fractions of interest, dispensing into aliquots, and flash freezing in N_2_ for storage at -80°C.

#### GFP-HEATR5B

Sf9 cells (2-L cell suspension at 2 x 10^6^ cells/mL) were infected with baculovirus incorporating the GFP-HEATR5B expression cassette and cultured for a further 72 h before cell pelleting by centrifugation. Cell pellets were lysed by resuspension and Dounce homogenisation (50 strokes) in lysis/washing buffer (LWB: 10 mM Tris-HCl pH 8.0, 150 mM NaCl, 0.5 mM EDTA, 1 mM DTT, 1X COMPLETE protease inhibitors and 2mM PMSF). The lysate was centrifuged for 30 min at 30,000 rpm using a Ti70 rotor. The supernatant was collected and loaded in a 5 mL Strep-Trap HP column (Cytiva) that was pre-equilibrated with LWB. After washing with 20 column volumns of LWB, the protein was eluted with ten column volumes of elution buffer (LWB including 3mM desthiobiotin (Sigma)). Fractions containing GFP-HEATR5B were pooled and the protein solution (3 mL) injected in a HiLoad 16/60 Superdex 200 Prep grade column (Cytiva) pre-equilibrated with gel filtration buffer (10 mM Tris HCl pH 7.4, 150 mM NaCl, 0.5 mM TCEP). Fractions were collected and analysed by SDS-PAGE, followed by pooling of relevant fractions, concentrating using an Amicon Ultra 4 centrifugal filter unit (100 kDa MWCO; Merck) to 1 mg/mL, dispensing into aliquots, and flash freezing in N_2_ for storage at -80°C.

#### Dynactin

Dynactin was purified from pig brain as described previously (Schlager *et al*., 2014; Urnavicius *et al*., 2015) and analysed by SDS-PAGE. Aliquots of the complex were flash frozen in N_2_ for storage at -80°C.

### Preparation of mouse brain extracts

Following dissection, brains from 16 C57BL/6j-OlaHsd male mice strain were transferred to bijou tubes containing solubilisation buffer (10 mM Hepes pH 7.3, 150 mM NaCl and 2 mM EDTA) on ice. Two brains were transferred to a pre-chilled 30-mL glass tube Wheaton homogenizer containing 5 mL of cold ‘solubilisation-plus’ buffer (solubilisation buffer with 1X COMPLETE protease inhibitor and 1X PhosSTOP phosphatase inhibitor (Roche)). The brains were homogenised in a coldroom using an electric homogeniser with a Teflon arm (1100-1300 rpm and 10 strokes). An extra 1 mL of solubilisation-plus buffer was added to the tube, followed by addition of two more brains and further homogenisation (15 strokes) with the same speed. The lysate was then transferred to a pre-chilled 50-mL falcon tube and kept on ice. The same procedure was repeated three times, followed by pooling of lysates from the 16 brains, addition of Triton X-100 (0.1% (v/v) final concentration) and incubating on ice for 15 min. The lysate was then split into eight pre-chilled, thick-walled polycarbonate tubes (3.2 mL, Beckman Coulter) and centrifuged in an ultracentrifuge (Beckman Optima TLX) using a TLA110 pre-chilled rotor at 70,000 rpm for 20 min at 4°C. The supernatants were pooled in a 50-mL falcon tube on ice and then flash frozen in N_2_ in 500 μL aliquots. Protein concentration of the extracts was ∼ 10 mg/mL.

### Pull-downs of dynein tail-associated proteins from brain extracts

#### Pull-downs using glutathione beads

Glutathione magnetic beads (Pierce) were washed twice with GST binding buffer (125 mM Tris-HCl pH 8.0, 150 mM NaCl, 0.05% Tween-20 and 1X COMPLETE protease inhibitor). 100 pmol (77 nM) of recombinant dynein tail complex or GST (Glutathione-S-transferase) only were incubated with 100 μL glutathione magnetic beads in a total volume of 1300 μL GST binding buffer. The beads were washed three times with 2 mL of GST binding buffer and incubated with 500 μL mouse brain extract (5 mg total protein) or 500 μL mouse brain extract spiked with 50 μg purified dynactin (41.67 pmol (48.45 nM)) in solubilisation-plus buffer for 2.5 h at 4°C with rotation. The beads were washed twice with 2 mL of Citomix buffer (60 mM KCl, 0.075 mM CaCl_2_, 5 mM K_2_HPO_4_/KH_2_PO_4_ pH 7.6, 12.5 mM HEPES pH 7.6, 2.5 mM MgCl_2_, 0.1% (v/v) Triton X-100, 2 mM ATP, 1X COMPLETE protease inhibitor and 1X PhosSTOP phosphatase inhibitor), twice with 50 mM ammonium bicarbonate pH 8.0, and resuspended in 100 μl of 50 mM ammonium bicarbonate pH 8.0. A 10-μL aliquot from each sample was saved for downstream quality control with immunoblotting by adding 10 μL 4X LDS buffer (Invitrogen) containing 200 mM DTT and incubating at 98°C for 15 min. The beads in these samples were magnetically separated for 10 min and the supernatant transferred to a fresh 1.5-mL tube that was then frozen at -20°C. The remaining 90 μL of bead slurry for each sample was processed for liquid chromatography-tandem mass spectrometry (LC-MS/MS), as described below.

#### Pull-down using IgG beads

IgG magnetic beads (Dynabeads M-280 Sheep α-Rabbit IgG, Life Technologies) were washed twice with IgG binding buffer (1X PBS, 0.05% (v/v) Tween-20 and 1X COMPLETE protease inhibitor). 100 pmol (77 nM) of recombinant dynein tail complex or recombinant protein A (ThermoFisher Scientific) were incubated with 100 μL IgG magnetic beads in 1300 μL IgG binding buffer with rotation for 1 h at 4°C. The beads were washed three times in IgG binding buffer and incubated with 500 μL mouse brain extract (5 mg total protein) or 500 μL mouse brain extract spiked with 50 μg purified dynactin (41.67 pmol (48.45 nM)) in solubilisation-plus buffer for 2.5 h at 4°C with rotation. The samples were washed and prepared for immunoblot analysis and LC-MS/MS following the same procedure described above for glutathione beads.

## LC-MS/MS

Proteins were digested on beads with 1 μg trypsin (Promega) for 18 h at 37°C, followed by acidification of peptides with 2% (v/v) formic acid. The bead/peptide mix was then centrifuged at 14,000 x g for 5 min and the supernatant collected. The peptide fractions (7 μL each) were analysed by nano-scale capillary LC-MS/MS with an Ultimate U3000 HPLC (ThermoScientific Dionex) with a 300 nL/min flow rate. Peptides were trapped in a C18 Acclaim PepMap 100 µ-precolumn cartridge (5 µm, 300 µm x 5mm (ThermoScientific Dionex)) prior to separation on a C18 Acclaim PepMap 100 (3 µm, 75 µm x 250 mm (ThermoScientific Dionex)) and elution with a 90-min gradient of acetonitrile (from 5% to 40%). Using a modified nano-flow electrospray ionisation source, the analytical column outlet was directly interfaced with a hybrid linear quadrupole ion trap mass spectrometer (Orbitrap QExactive (ThermoScientific)). Data-dependent analysis was carried out with a resolution of 60,000 for the full MS spectrum (collected over a 200–1800 m/z range), followed by ten MS/MS spectra in the linear ion trap (collected using 35-threshold energy for collision-induced dissociation).

### Mass-spectrometry data processing and analysis

Raw mass-spectrometry data from pull-down samples were processed with MaxQuant software (versions 1.5.6.2; (Tyanova *et al*., 2016a)) using the built-in Andromeda engine to search against the UniprotKB mouse proteome (Mus musculus; release 2012_02) containing forward and reverse sequences. The iBAQ algorithm and “Match Between Runs” option were additionally used. Carbamidomethylation was set as a fixed modification, and methionine oxidation and N-acetylation were set as variable modifications (using an initial mass tolerance of 6 ppm for the precursor ion and 0.5 Da for the fragment ions). For peptide and protein identifications, search results were filtered with a false discovery rate (FDR) of 0.01. Datasets were further processed with Perseus software (version 1.6.13.0; (Tyanova *et al*., 2016b)). Protein tables were filtered to eliminate identifications from the reverse database, as well as common contaminants. Only proteins identified on the basis of at least two peptides and a minimum of three quantification events in at least one experimental group were taken forward. iBAQ intensity values were normalised against the median intensity of each sample (using only those peptides that had intensity values recorded across all samples and biological replicates), followed by log_2_-transformation and filling of missing values by imputation with random numbers drawn from a normal distribution calculated for each sample, as previously described (Neufeldt *et al*., 2019; Plaszczyca *et al*., 2019). Proteins that were statistically significantly enriched between pairs of datasets were identified with Welch’s t-tests with permutation-based false discovery rate statistics. We performed 250 permutations and the FDR threshold was set at 0.05. The parameter S0 was set at 0.1 to separate background from specifically enriched interactors. Volcano plots of results were generated in Perseus. UniprotKB accession codes of all protein groups and proteins identified by mass spectrometry are provided in Supplementary datasets 1 – 4.

### Flp-In T-REx cell line generation

HEK293 or HeLa Flp-In cells in wells of 6-well plates (8.5 x 10^5^ cells/well) were transfected with 1 μg plasmid DNA (pcDNA5 FRT/TO GFP-linker or pcDNA5 FRT/TO GFP-HR5B with pOG44 in a 1:1 or 2:1 ratio, respectively). The DNA was mixed with 200 μL OptiMEM (Gibco) and 5 μl 1mg/mL Polyethyleneimine (PEI) “MAX” MW 40,000 (Polysciences) in sterile phosphate-buffered saline (PBS). After vortexing, the mixture was incubated for 15 min at 20°C before adding to the cells in a drop-wise manner and gentle swirling of the plate. Cells were then incubated for 24 h at 37°C before removal of the media and splitting into a T25 flask containing 150 μg/mL hygromycin B and 5 μg/mL blasticidin (both from Gibco). The next day, and once every following week, the selective media was changed until single colonies appeared, which were detached with trypsin and pooled together for subsequent experimentation.

### GFP-Trap immunoprecipitation from extracts

To induce expression of GFP or GFP-HR5B, HEK293 or HeLa Flp-In T-REx GFP-linker or GFP-HR5B cell lines cells were cultured in the presence of 1 μg/mL tetracycline for at least 48 h. Cells from three 15-cm dishes were harvested for each immunoprecipitation sample and lysed in 1.2 mL of ice-cold lysis buffer (10mM Tris-HCl pH 7.4, 150 mM NaCl, 0.5mM EDTA, 5% NP-40, 1 mM PMSF, 1X COMPLETE protease inhibitors and 1X PhosSTOP phosphatase inhibitors) for 30 min by pipetting extensively every 10 min. Lysates were transferred to pre-chilled thick polycarbonate tubes and centrifuged at 30,000 rpm for 30 min in a TL110 rotor using a Beckman Optima TLX ultracentrifuge. The supernatant was transferred to a pre-cooled 15-mL falcon tube, followed by addition of 1.5 volumes of ice-cold dilution buffer (10 mM Tris/HCl pH 7.4, 150 mM NaCl, 0.5mM EDTA, 1X COMPLETE protease inhibitor and 1X PhosSTOP phosphatase inhibitor). The diluted supernatant was added to 60 μL of equilibrated GFP-Trap®_MA bead slurry (Chromotek) and the samples tumbled end-over-end for 4 h at 4°C. Beads were washed in 1 mL Citomix buffer (with mixing by pipetting up and down four times), transferred to fresh tubes and denatured with LDS/DTT as described above for pull-downs from mouse brain extracts.

### *In vitro* protein-protein interaction assay

For each sample, 25 μL of GFP-Trap®_MA bead slurry (Chromotek) was equilibriated with 1 mL of cold dilution buffer (DB: 10 mM Tris-HCl pH 7.4, 150 mM NaCl, 0.5 mM EDTA, 1X COMPLETE protease inhibitors and 1X PhosSTOP phosphatase inhibitors), followed by blocking of non-specific binding sites on the beads with 4% bovine serum albumen (BSA) in DB for 50 min at 4°C with end-over-end tumbling. Beads were washed twice in DB and transferred to fresh 1.5-mL tubes. Following removal of the supernatant, 80 pmol of purified GFP or GFP-HR5B in 0.5 mL of DB containing 2% NP-40 and 0.4 mM PMSF was mixed with the beads by end-over-end tumbling at 4°C for 2 h. Beads were washed three times in RIPA buffer (10 mM Tris-HCl pH 7.4, 150 mM NaCl, 0.5 mM EDTA, 0.1% SDS, 1% Triton X-100, 1 X COMPLETE protease inhibitors and 1X PhosSTOP phosphatase inhibitors) and two times in protein binding buffer (PBB: 10mM HEPES pH 7.6, 150 mM NaCl, 0.5 mM EDTA, 0.1% Triton X-100, 1% deoxycholate and 1X COMPLETE protease inhibitors), followed by transfer to fresh tubes and removal of the supernatant. Dynein tail (20 pmol), dynactin (10 pmol) or dynein tail with dynactin (20 pmol and 10 pmol, respectively) were incubated with the beads in 0.5 mL PBB for 2 h at 4°C with end-over-end tumbling. Subsequently, beads were washed twice in 1 mL RIPA buffer (with mixing by pipetting up and down four times), transferred to a fresh tube and processed for immunoblotting as described above. One third of the sample was loaded per gel lane, alongside 0.1 μg of dynein tail or dynactin alone for molecular weight comparison.

### Immunoblotting

Proteins were separated on NuPAGE Bis-Tris gels (4% – 12%) (ThermoFisher Scientific), with ECL Rainbow Full Range Marker (Cytiva) used for molecular weight standards. Following transfer to methanol-activated PDVF membrane (Immobilon P, Millipore) using the XCell II blot system (ThermoFisher Scientific), membranes were blocked with 5% (w/v) milk powder (Marvel) in PBS, and incubated with primary antibodies in 1% milk powder or 1% BSA in PBS overnight at 4°C. After washing with PBS/0.5% Tween-20 or PBS/1% Tween-20, membranes were incubated with secondary antibodies in 1% milk powder or 1% BSA in PBS for 1 h at room temperature. Details of primary and antibodies are provided below. Signals were developed using the ECL Prime system (Cytiva) and Super RX-N medical X-ray film (FUJIFILM). When loading samples for immunoprecipitation experiments, the volume of extract samples (input) was adjusted to give a clear, non-saturated signal (5 μL for AP1γ and

γ-SYNRG, 20 μL for AFTPH, 2 μL of a 1/20 dilution for DYNC1H1 and GFP, 20 μL of a 1/20 dilution for GFP-HR5B, 8 μL of a 1/20 dilution for DCTN1 and BICD2 and 4 μL of a 1/20 dilution for GAPDH). For all immunoprecipitates, one third of the captured sample was loaded per gel lane.

### siRNA transfection

The *DYNC1H1* siRNA (ON-TARGET plus *DYNC1H1* L-006828-00-0005) and non-targeting control siRNA (ON-TARGET plus D-001810-10-05) pools were synthesised by Dharmacon and resuspended in 1X siRNA buffer (Dharmacon). 2.5 x 10^4^ U2OS cells/well were seeded in 24-well plates a day before transfection. 1.5 μL Lipofectamine RNAiMax reagent (Invitrogen) was added to 50 μL OptiMEM (Gibco) containing 25 nM siRNA and the mix incubated for 15 – 20 min at room temperature. This transfection mix was then added dropwise to the culture medium (450 μL/well), which had been replaced 30 min earlier, and mixed by gently shaking the plate. 48 h after transfection, cells were washed with PBS and fixed with 4% paraformaldehyde (Sigma)/PBS for 20 min at room temperature. After three washes with PBS, cells were processed for immunofluorescence (see below).

### Transient transfection of human cells

U2OS cells were reverse transfected with pcDNA3.1-eGFP-HR5B (400ng/well) or pcDNA3.1-GFP linker (200ng/well), which was pre-mixed with 50 μL OptiMEM and 1.25 μL FuGene (Promega) for 15 – 20 min at room temperature. The transfection mix was added on coverslips, followed by addition of 7 x 10^4^ cells in 120 μL complete DMEM and adjustment of the final volume to 500 μL using additional complete DMEM. 24 h later, the transfection medium was removed and cells fixed in 4% PFA, washed in PBS and processed for immunofluorescence (see below).

### Generation of HEATR5B deficient U2OS cell lines

Negative control scrambled sgRNA (variable sequence: GCACUACCAGAGCUAACUCA) and *HEATR5B* +37079270 (reverse) sgRNA (variable sequence: GGAUUAAUAAGUAGUUCACC) (CRISPRevolution; Synthego) were resuspended in nuclease-free 1X TE buffer (10 mM Tris-HCl, 1 mM EDTA pH 8.0). U2OS cells were electroporated with ribonucleoprotein complexes containing sgRNA and Cas9 2NLS protein (Synthego) using the ThermoFisher Scientific Neon^TM^ transfection system according to the manufacturer’s instructions (2.5 x 10^5^ cells in 7 μL buffer R (ThermoFisher Scientific) and 7 μL sgRNA:Cas9 RNP mix (90 pmol:10 pmol)). Cells were then seeded in wells of a 6-well plate containing 4 mL pre-warmed complete DMEM. Six days later, limited dilution of cells in 96-well plates was performed to isolate single clones. Colonies of these clones were harvested by trypsin treatment and expanded for DNA extraction. The target region in the *HEATR5B* gene was amplified from genomic DNA according to Synthego’s protocol and analysed by Sanger sequencing. Primers F-KO HEATR5B (TGG CTT TGG AGG AGC ATG AAG) and R-KO HEATR5B (ACT TCA AGG GCC CCT ATT AAA G) were used for PCR, with primer SP HEATR5B (GAG TGC CTT AAG TGT TAA GTG TTT) used for sequencing of the product. Indels were identified in sequencing chromatograms using the ICE application (Inference of CRISPR Edits; Synthego).

### Antibodies for immunoblotting and immunofluorescence of human cells

The following antibodies were used for immunoblotting (IB) and immunofluorescence (IF) of human cell extracts or cells, respectively (working dilutions in parentheses): α-GFP chicken antibody (Abcam 13970; IB, 1:5000; IF, 1:200), α-DYNC1H1 rabbit antibody (Proteintech 12345-1-AP; IB, 1:100; IF, 1:100), α-DCTN1 mouse antibody (BD Transduction Laboratories 610474; IB, 1:5000; IF, 1:50), α-AFTPH rabbit antibody (Invitrogen PA5-57104; IB, 1:500), α-γ-SYNRG (AP1GBP1) rabbit antibody (Novusbio NBP1-90145; IB, 1:200), α-AP1γ (F-10) mouse antibody (Santa Cruz Sc-398867; IB, 1:100), α-AP1γ mouse antibody (Sigma A4200, clone 100.3; IF, 1:400), α-EEA1 mouse antibody (BD Transduction laboratories 610457; IF, 1:200), α-LAMP1 mouse antibody (Abcam 25630; IF, 1:200), α-GAPDH rabbit antibody (Sigma G9545; IB, 1:4000), α-HEATR5B rabbit antibody ((Hirst *et al*., 2005); provided by J. Hirst and M. Robinson (Cambridge Institute for Medical Research, UK); IB, 1:250), α-TGN46 sheep antibody (Bio-Rad AHP500; IF, 1:200), α-RAB11A rabbit monoclonal antibody (Abcam 128913); IF, 1:50), α-β-actin mouse antibody (Gen Tex GTX26276; IB, 1:20,000), α-α-Tubulin mouse antibody (Santa Cruz; Sc-32293 clone DM1A; IF, 1:500), and α-Transferrin receptor rabbit antibody (Abcam ab84036; IF, 1:200).

### Immunofluorescence of human cells

Control and *HEATR5B* mutant cells, GFP or GFP-HEATR5B transfected cells, or GFP and GFP-HEATR5B Flp-in cells (cultured in the presence of tetracycline for 72 h) were fixed with 4% PFA for 20 min at room temperature. After three washes with PBS, cells were permeabilised with 0.1% Triton X-100 in PBS for 10 min, blocked with either 10% FBS (Gibco) or 1% BSA in PBS for 30 min and incubated with primary antibodies in blocking solution (1% FBS/PBS or 1% BSA/PBS) for 1 h at room temperature. After washing twice with PBS, secondary antibodies (Alexa-conjugated IgG H+L series, ThermoFisher Scientific; 1:1000 dilution) were incubated with cells for 1 h. Cells were washed twice more in PBS and incubated in 1 μg/mL DAPI (Sigma) in PBS for 2 min to stain DNA, followed by two further PBS washes. Cells were washed once with autoclaved MilliQ water before mounting using ProLong Gold anti-fade mountant (ThermoFisher Scientific). Imaging was performed on a Zeiss 710 or 780 confocal microscope, with laser and acquisition settings kept constant within an experimental series.

### Live imaging of human cells and mean square displacement analysis

Cells were seeded on 35-mm Fluorodish dishes (Fisher Scientific) in 2 mL complete DMEM. An hour before imaging, cell medium was replaced with pre-warmed Leibovitz medium without phenol red (Gibco), which was supplemented with 20 mM HEPES and 10% FBS. For imaging of AP1σ1-RFP or DsRed-RAB11A in GFP-HR5B cells, as well as AP1σ1-RFP in control and *HR5B* KO cells, 1 x 10^5^ cells were seeded the day before transfection and cell culture medium replaced with 1.5 mL fresh medium (without tetracycline) 1 h before transfection. Cells were transfected with a transfection mix comprising 500 ng pAP1σ1-RFP or 400 ng pDsRed-RAB11A WT in 250 μL OptiMEM and 6.25 μL PEI that had been pre-incubated at room temperature for 20 min before transfection. The transfection medium was replaced 6 h post transfection with fresh pre-warmed medium (containing tetracycline). Transfected cells showing low-to-medium expression of AP1σ1-RFP or DsRed-RAB11A were filmed 24 h post transfection.

Movies of GFP-HR5B cells expressing DsRed-RAB11A were acquired using the 63 X 1.4 NA Plan Apo oil objective of a PerkinElmer Ultraview ERS confocal spinning disk system built around an Olympus IX71 microscope equipped with an Orca ER camera (Hamamatsu). Data were acquired using sequential imaging of the two channels (one pair of images captured every 1.3 s). Movies of GFP-HR5B cells expressing AP1σ1-RFP were acquired on a custom spinning disk confocal microscopy system composed of a Nikon Ti stand equipped with perfect focus, a fast-piezo z-stage (ASI) and a 100X NA 1.49 TIRF objective. Confocal illumination was achieved with a CSU-X1 spinning disk head (Yokogawa) and a Photometrics 95B back-illuminated sCMOS camera operating in global shutter mode and synchronised with the rotation of the spinning disk. Excitation was performed with 488-nm (150 mW OBIS LS) and 561-nm (100 mW OBIS LS) lasers fibered within a Cairn laser launch. To enable fast acquisition, hardware was synchronised by a Zynq-7020 Field Programmable Gate Array (FPGA) stand-alone card (National Instrument sbrio 9637) running custom code. Sample temperature was maintained at 37°C using a heating enclosure (MicroscopeHeaters.com). The camera was configured in 12-bit dynamic range mode (gain 1) to maximise sensitivity, and a 600 x 600 pixel region of interest selected to maximise acquisition speed. Sequential imaging of a single z-plane was performed with a total exposure time for both frames of 0.206 s. Absolute timestamps were recorded to ensure accurate tracking. Acquisition was controlled by Metamorph software (7.10.1.161).

High-speed imaging of AP1ο−1-RFP motility in human cells was performed using the Nikon Ti-based custom spinning disk confocal microscopy system described above, with the exception that the camera was operated in streaming mode with an effective frame-rate of 17.5 Hz (50 ms exposure plus ∼7 ms read-out time). Particle tracking and analysis of these time series was performed in Fiji (Schindelin *et al*., 2012) and Matlab 2021b (Mathworks). AP1ο−1-RFP particles were automatically detected in a threshold-free manner by 2D gaussian fitting using the TwoTone TIRF-FRET matlab package (Holden *et al*., 2010). Trajectories were then automatically tracked using a MATLAB implementation by D. Blair and E. Dufresne of the IDL particle tracking code initially developed by D. Grier, J. Crocker and E. Weeks (http://site.physics.georgetown.edu/matlab/index.html). Since short tracks make MSD analysis inaccurate (Zahid *et al*., 2018), tracks that lasted for fewer than 10 time points were discarded. We also manually drew Regions of Interest (ROI) around cells and excluded all tracks corresponding to tracked objects that were not located in cells. For each track, the MATLAB class MSD Analyzer (Tarantino *et al*., 2014) was used to compute the Mean Square Displacement (MSD) of segments of increasing duration (delay time *t*) (*MSD* (*t*) = < (Δ*x*)^2^ > + < (Δ*y*)^2^ >). The weighted mean MSD (Tarantino *et al*., 2014) across all tracks was then computed per delay time and plotted as a function of the delay time considered. AP1ο−1-RFP particle intensity upon *HR5B* disruption or as a function of track length was computed by integrating the area under the gaussian from the gaussian fitting data, followed by averaging of the values per track.

### *Drosophila* culture and preexisting strains

*Drosophila melanogaster* strains were cultured using ‘Iberian’ food (5.5% (w/v) glucose, 3.5% (w/v) organic wheat flour, 5% (w/v) baker’s yeast, 0.75% (w/v) agar, 16.4 mM methyl-4-hydroxybenzoate (Nipagin), 0.004% (v/v) propionic acid). Stocks and crosses were maintained in an environmentally controlled room set to 25 ± 1°C and 50 ± 5% relative humidity with a repeating 12h-light/12h-dark regime. The following previously generated strains were used in the study: *nos-Cas9^ZH-2A^* (Bloomington *Drosophila* Stock Center stock number: BL54591; (Port *et al*., 2014)), *Df(3R)BSC222* (BL9699; containing a genomic deficiency that includes the *Heatr5* locus); and *YFP-Rab11* (Dunst *et al*., 2015). Wild-type flies were of the *w^1118^* strain.

### Generation of *Drosophila Heatr5* mutant, GFP-Heatr5B and GFP-Golgin-245 strains

Flies expressing a pair of gRNAs that target the 5’ region of the *Heatr5* gene (gRNAs 1+2; Figure S8A) from the U6:3 promoter were generated using the pCFD4 plasmid, as described (Port *et al*., 2014) (see Table S4 for sequences of oligos used for gRNA cloning). Males expressing this transgene were crossed with *nos-Cas9^ZH-2A^* females (which express Cas9 specifically in the germline), with mutations identified in the offspring of the progeny by Sanger sequencing of PCR products derived from the targeted genomic region (Port & Bullock, 2016). The GFP-Heatr5 expression construct was generated by restriction enzyme-mediated cloning of PCR-amplified eGFP and Heatr5 coding sequences into a pCASPER-based plasmid that expresses proteins under the control of the ubiquitously active α-tubulin84B promoter (Dienstbier *et al*., 2009). The Heatr5 sequence was derived from the full-length cDNA clone GH08786 (*Drosophila* Genome Resource Center). Flies were transformed with this plasmid by P-element-mediated integration using standard procedures. The GFP-Golgin-245 knock-in strain was generated by CRISPR/Cas9-mediated homology-directed repair (HDR), as described (Port & Bullock, 2016; Port *et al*., 2014). Briefly, embryos of *nos-Cas9^ZH-2A^* mothers were injected with a mixture of three plasmids: two pCFD3-gRNA plasmids targeting the *Golgin245* genomic sequences shown in Figure S12 (see Table S4 for oligo sequences used for cloning) and a pBlueScript-based donor plasmid (pBS-GFPG245) that has eGFP flanked by ∼1-kb *Golgin245* homology arms. The final concentration of each plasmid in the injection mix was 100 ng/μL for each of pCDF3-gRNA plasmids and 150 ng/μL for pBS-GFPG245. Successful HDR was confirmed by PCR-based analysis of the progeny of the offspring of surviving embryos, as described (Port & Bullock, 2016; Port *et al*., 2014; Port *et al*., 2015).

### CRISPR-based disruption of *Heatr5* in the female *Drosophila* germline

Mothers doubly heterozygous for *nos-Cas9^ZH-2A^* and one of two pCDF4 transgenes that express independent pairs of gRNAs targeting Heatr5 (gRNAs 1+2 or gRNAs 3+4; Figure S8A and S9A) were crossed with wild-type males and introduced into egg-laying cages mounted on plates of apple juice agar (1.66% (w/v) sucrose, 33.33% (v/v) apple juice, 3.33% (w/v) agar, and 10.8 mM methyl-4-hydroxybenzoate). Embryos from this cross were collected in timed egg lays and processed for immunofluorescence, microinjection or immunoblotting as described below.

### Immunostaining and cuticle preparations of *Drosophila* embryos

For immunostaining, embryos were dechorionated, washed, fixed with a 4% formaldehyde/n-heptane mixture, and devitellinised using standard procedures. Washes were performed in PBS/0.1% Triton X-100 (PBST) and embryos blocked in 20% Western Blocking Buffer (Sigma) in PBST. The following primary antibodies were used: α-*Drosophila* AP1γ rabbit antibody ((Hirst *et al*., 2009); provided by J. Hirst and M. Robinson (Cambridge Institute for Medical Research, UK; 1:1000 dilution); α-GFP chicken antibody (Abcam ab13970; 1:250 dilution); α-*Drosophila* GM130 rabbit antibody (Abcam 30637; 1:1000 dilution) and α-*Drosophila* Golgin-84 mouse antibody ((Riedel *et al*., 2016); provided by S. Munro (MRC-LMB, Cambridge, UK; 1:50 dilution)). Secondary antibodies were from the Alexa-conjugated IgG H+L series (ThermoFisher Scientific; 1:500 dilution). Samples were mounted in Vectashield containing DAPI (Vector Laboratories) and imaged with a Zeiss 780 laser-scanning confocal microscope. For cuticle preparations, 0 – 4 h egg collections were incubated for a further 28 h at 25°C. Unhatched embryos were then dechorionated, washed and mounted in lacto-hoyers solution before clearing by heating in an oven at 65°C for 16 h.

### Analysis of immunofluorescence signals in human cells and fly embryos

Quantification of spread of fluorescent signals away from the nucleus in U2OS cells was performed using a custom ImageJ script (available at https://github.com/jboulanger/imagej-macro/tree/main/Cell_Organization). The nuclear DAPI signal was segmented either using StartDist (Schmidt *et al*., 2018) or a median filter, followed by a rolling ball, a threshold controlled by a probability of false alarm, a watershed and the extraction of connected components. The signed euclidean distance transform was then computed for each region of interest, which allowed association of each pixel with the closest nucleus. Background pixels were discarded by segmenting the maximum projection intensity of all channels using a smoothing step and a threshold based on a quantile of the intensity distribution. Regions touching the border of the image were discarded to avoid analysis of incomplete cells. The signal of interest was segmented using a sharpening, a median filter, a rolling ball and finally a threshold controlled by a user defined false alarm rate. For each cell, the spread was computed as the sum of the squared deviation to the centroid weighted by the signal intensity in each segmented mask. This procedure was implemented as a macro to process batches of images. The number and length of tubules was determined manually in ImageJ using the freehand selection tool. ImageJ was also used for quantification of particle number and total particle intensity in mammalian cells and fly embryos. The non-adjusted channel of interest was thresholded and segmented, followed by removal of particles smaller than five pixels. In mammalian cells, particle intensity and number of particles were measured per individual cell.

In fly embryos, rectangular regions of 300 μm^2^ were analysed in the apical cytoplasm, upper basal cytoplasm, lower basal cytoplasm and yolk; the upper basal and lower basal cytoplasmic regions were defined by the midpoint of the distance between the edge of the yolk and the basal side of the nuclei.

### Analysis of AP1 and mRNA motility in *Drosophila* embryos

Motility of AP1-positive structures was assessed with an affinity purified α-*Drosophila* AP1γ rabbit antibody that has been shown to be highly specific (Hirst *et al*., 2009). Fluorescent AP1γ antibody conjugates were generated by incubating 1 μL of the α-AP1γ antibody with 0.5 μL of Alexa555-conjugated donkey α-rabbit IgG (H+L) highly cross-adsorbed secondary antibody (ThermoFisher Scientific) and 3.5 μL PBS at room temperature for 30 min. The antibody solution was then dialysed by dropping on a 0.025 μm-pore filter MCE membrane (13 mm-diameter; Millipore) that was floating on 25 mL of PBS, and incubating for 1 h. The drop was recovered and centrifuged briefly to remove any aggregates before loading into a laser-pulled borosilicate Microcap needle (Drummond). Primary antibodies conjugated to the secondary antibody as negative controls were: rabbit α-HA (Sigma H6908) and rabbit α-dinitrophenol (Invitrogen A6430). Alexa488-UTP-labelled *bcd* mRNA was generated by *in vitro* transcription as described (Bullock *et al*., 2006) and loaded into the injection needle at a concentration of 750 ng/μL in dH_2_O.

Antibody and mRNA injections into embryos were performed at room temperature as described (Bullock *et al*., 2006) using a micromanipulator fitted to an Ultraview ERS spinning disk imaging system (PerkinElmer). Time-lapse images were acquired at 1 frame/s (AP1γ antibody conjugates) or 2 frame/s (*bcd* mRNA) using a 60×/1.2 NA UPlanApo water objective and an OrcaER camera (Hamamatsu). Centroid-based automatic tracking of fluorescent antibody and mRNA particles in the cytoplasm and quantification of their movements was performed using a custom script written in Mathematica (Wolfram) by A. Nicol and D. Zicha (Bullock *et al*., 2006). To assess the contribution of microtubules to motility of AP1-positive puncta, wild-type embryos were injected with a 20 ng/μL solution of colcemid 2 min prior to injection of the fluorescent AP1γ antibody conjugate. After imaging the antibody signal for ∼10 min, colcemid was inactivated by a 10 s exposure to UV light through the mercury lamp attached to the microscope, with filming resumed 60 s afterwards. To determine the role of dynein in transport of AP1 positive structures, embryos were preinjected 2 min before AP1γ antibody conjugate injection with a 1 μg/μL solution of mouse α-Dic antibody (clone 74.1; Millipore) or a 1 μg/μL solution of mouse α-GFP antibody (mixture of clones 7.1 and 13.1; Sigma) as a control. These antibodies were dialysed before use, as described above. Multiple injection sessions were performed for each experiment to confirm the consistency of results. When assessing the effects of mutating *Heatr5*, the control and mutant embryo injections were interleaved within each imaging session.

To visualise both Golgin-245 and AP1 in the live blastoderm, embryos from GFP-Golgin-245 homozygous mothers were injected with Alexa555-labelled AP1γ antibody conjugates as described above. To maximise the intensity of the relatively dim GFP-Golgin-245 signal, the focal plane was set closer to the surface of the embryo than for AP1γ or mRNA tracking. This meant that it was typically not possible to follow transported AP1γ puncta all the way from the yolk to the upper basal cytoplasm. In these experiments, images were subjected to 2 x 2 binning, with sequential imaging of the two channels (exposure times of 500 ms for the AP1γ signal and 1 s for the GFP-Golgin-245 signal).

### Immunoblotting of *Drosophila* embryo extracts

*Drosophila* embryo extracts were generated for immunoblotting as described (McClintock *et al*., 2018), with a Coomassie (Bradford) Protein Assay (ThermoFisher Scientific) used to ensure that the amount of total protein loaded per gel lane was equivalent between genotypes. Electrophoresis and immunoblotting were performed as described above using α-*Drosophila* AP1γ rabbit antibodies (see above; 1:5000 dilution) and α-β-actin rabbit antibodies (Abcam 8224; 1:10,000 dilution).

### Statistical analysis and data plotting

Statistical evaluations and plots were made with Prism (version 9; Graphpad) or Perseus. Details of sample sizes and types of tests are included in the figure legends. Individual data points are shown in plots, except when N > 40 (when box and whisker plots are used).

## DATA AVAILABILITY

The mass spectrometry-based proteomics data will be deposited at the ProteomeXchange Consortium (http://proteomecentral.proteomexchange.org) via the PRIDE partner repository.

## AUTHOR CONTRIBUTIONS

V.M. and S.L.B. conceived the project. V.M., L.A.A., L.J., J.L.W., N.M., F.B., K.B. and S.L.B. devised and performed experiments, V.M., L.A.A., L.J., J.B., N.M., J.L.W., P.S., E.D. and

S.L.B. analysed and interpreted data. V.M., E.D., M.A.K. and S.L.B. supervised experiments.

S.L.B. co-ordinated the project. V.M. and S.L.B. prepared the manuscript, which was edited and approved by each of the other authors.

## COMPETING INTERESTS STATEMENT

The authors have no competing financial interests.

## ACKNOWLEDGEMENTS

We are very grateful to David Paul, Jennifer Hirst, Harvey McMahon, Alfredo Castello, Elena Dobrikova, Matthias Gromeier, Eeson Rajendra, Alice Bittleston, Sabine Thomas, Andrew Carter and members of the Bullock and Carter groups (in particular Mark McClintock and Sami Chaaban) at MRC-LMB for discussions, advice, assistance or reagents, Daniel Zicha for help with analysis of particle tracks in *Drosophila* embryos, Mark Skehel for helping process mass spectrometry data, Jo Westmoreland for contributing artwork, Graham Lingley for help with editing movies, and Sean Munro and Margaret Robinson for reagents, advice and comments on the manuscript. This work was supported by UK Research and Innovation (MRC file reference numbers MC_U105178790 (to S.L.B.), MC_UP_1201/13 (to E.D.) and MC_U105178783 (for support of N.M. (award to Sean Munro)), as well as the Human Frontier Science Program (Career Development Award CDA00034/2017-C to E.D). Work in the group of P.S. is supported by the Free and Hanseatic City of Hamburg and the Federal Ministry of Education and Research (BMBF, project VirMScan). Work in the group of M.A.K. is supported by the DFG (Ki 502/9-1, 506658941), as well as WifoMed and the Friedrich-Baur-Stiftung (awards to K.B). For the purpose of open access, the MRC Laboratory of Molecular Biology has applied a CC-BY public copyright licence to any Author Accepted Manuscript version arising.

## SUPPLEMENTARY FIGURES

## SUPPLEMENTARY DATASET LEGENDS

**Supplementary dataset 1. Mass spectrometry-derived iBAQ values for dynein tail-GST and GST only pulldowns when exogenous dynactin was absent from both conditions.** In these and other datasets, the first tab shows the complete dataset and the second tab shows statistically significantly enriched proteins in the dynein-tail vs control pulldown (q value <0.05). For calling of hits (highlighted in pink), an additional threshold of iBAQ value fold change for dynein tail vs control of >10 was applied. In tab 2, proteins that are not core components of the dynein or dynactin complex are labelled in bold.

**Supplementary dataset 2. Mass spectrometry-derived iBAQ values for dynein tail-GST and GST only pulldowns when exogenous dynactin was present in both conditions.**

**Supplementary dataset 3. Mass spectrometry-derived iBAQ values for ZZ-dynein tail and protein A only pulldowns when exogenous dynactin was absent from both conditions.**

**Supplementary dataset 4. Mass spectrometry-derived iBAQ values for ZZ-dynein tail and protein A only pulldowns when exogenous dynactin was present in both conditions.**

## SUPPLEMENTARY MOVIE LEGENDS

**Movie S1. Dual colour movie of HeLa cell expressing GFP-HEATR5B and AP1σ1-RFP** Each loop corresponds to 13 s of real time.

**Movie S2. Example of long-distance co-transport of GFP-HEATR5B and AP1σ1-RFP in HeLa cell cytoplasm (crop of time series used to produce Movie S1).** Shown is a composite for individual channels and the merge. Yellow arrow shows particle that will undergo long-distance movement. Nucleus is positioned to the bottom right of the frames. Movie corresponds to 19 s of real time.

**Movie S3. Dual colour movie of HeLa cell expressing GFP-HEATR5B and dsRed-RAB11A.** Each loop corresponds to 117 s of real time.

**Movie S4. Example of long-distance co-transport of GFP-HEATR5B and dsRed-RAB11A in HeLa cell cytoplasm (crop of time series used to produce Movie S3).** Shown is a composite of individual channels and the merge. Yellow arrow shows particle that will undergo long-distance movement. Nucleus is positioned at the bottom of the frames. Movie corresponds to 79 s of real time.

**Movie S5. Behaviour of AP1σ1-RFP in control and *HEATR5B* KO U2OS cells.** Image series from movies with overlaid tracks. Time is shown in the bar in the lower right corner of the composite. Note that the control movie has two cells in the field of view.

**Movie S6. Behaviour of injected Alexa555-secondary antibody/AP1γ primary antibody conjugates in control (*nos-cas9*) and *nos-cas9 gRNA-Hr5^1+2^* blastoderm embryos.** Apical is to the top. Movie corresponds to 252 s of real time.

**Movie S7. Behaviour of injected Alexa555-secondary antibody/AP1γ primary antibody conjugates in wild-type embryo preinjected with colcemid and after inactivation of colcemid with UV light.** ‘UV’ indicates period of exposure to UV light (1 min 5 sec into the duration of the movie). Movie corresponds to 600 s of real time before colcemid inactivation with a 10 s pulse of UV and 375 s of real time after inactivation. Movie is representative of injections into four embryos.

**Movie S8. Behaviour of injected Alexa555-secondary antibody/AP1γ primary antibody conjugates in wild-type embryo preinjected with α-GFP or α-Dic antibodies.** Apical is to the top. Movie corresponds to 402 s of real time.

**Movie S9. Behaviour of injected Alexa488-labelled *bcd* RNA in control (*nos-cas9*) and *cas9 gRNA-Hr5^1+2^* blastoderm embryos.** Apical is to the top. Movie corresponds to 430 s of real time.

**Movie S10. Behaviour of injected Alexa555-secondary antibody/AP1γ primary antibody conjugates with respect to GFP-Golgin-245.** Apical is to the top. Squares show regions and time periods in Movie S11 and S12. Movie corresponds to 879 s of real time.

**Movie S11. Example of interaction of transported AP1γ particle with GFP-Golgin-245 in the upper region of the basal cytoplasm (crop of Movie S11).** Yellow arrow shows AP1γ punctum that is transported apically, interacts with the Golgi, and is transported basally. Apical is to the top. Each loop corresponds to 36.1 s of real time.

**Movie S12. Additional example of co-transport of AP1γ and GFP-Golgin-245 in the upper region of the basal cytoplasm (crop of Movie S11).** Yellow arrow shows an example of bidirectional co-transport of AP1γ in association with Golgin-245. Apical is to the top. Each loop corresponds to 56 s of real time.

## SUPPLEMENTARY TABLES

**Figure S1.**
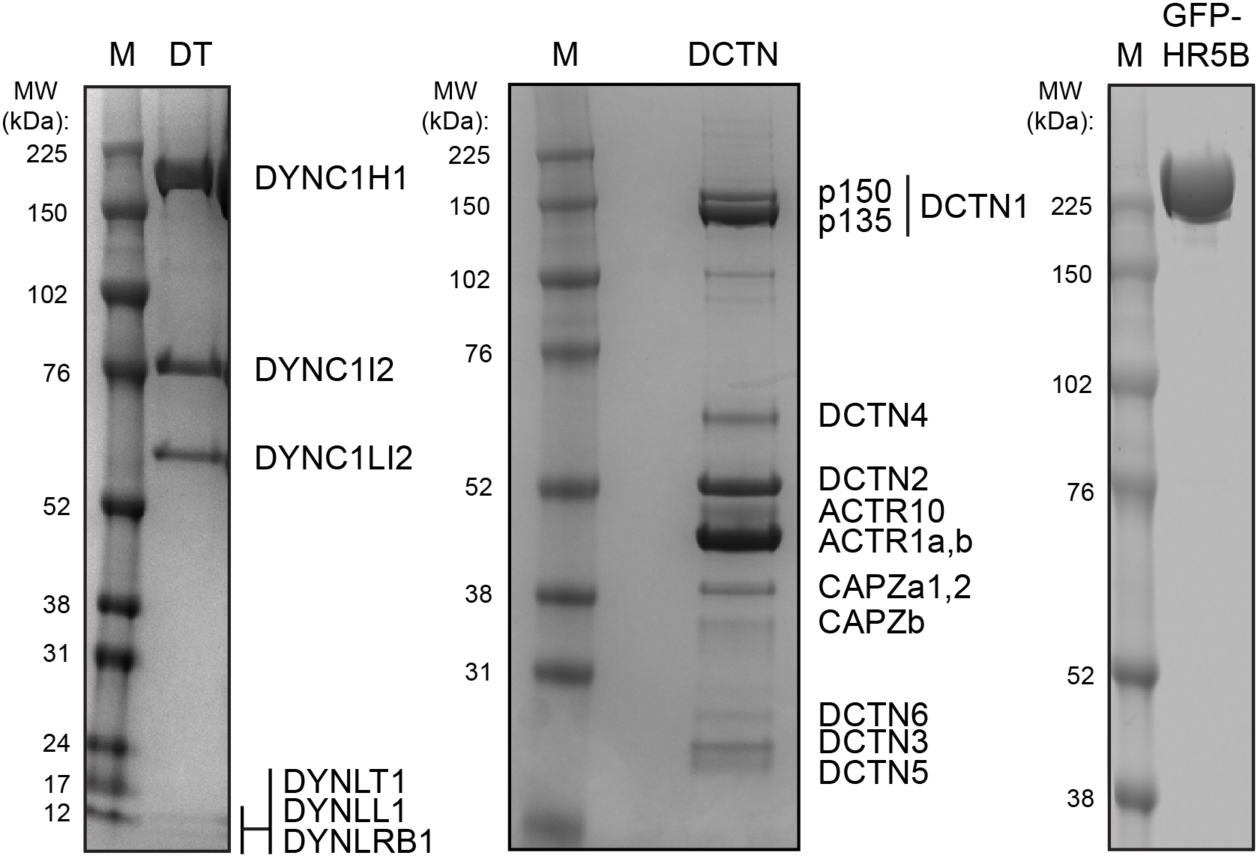
Purified protein samples. Images of Coomassie-stained gel lanes after electrophoresis of the human dynein tail complex (DT), pig brain dynactin (DCTN) and GFP tagged human HEATR5B (HR5B). Note that DCTN1 exists as two different isoforms (p150 and p135). M, protein markers; MW, molecular weight of protein markers.

**Figure S2.**
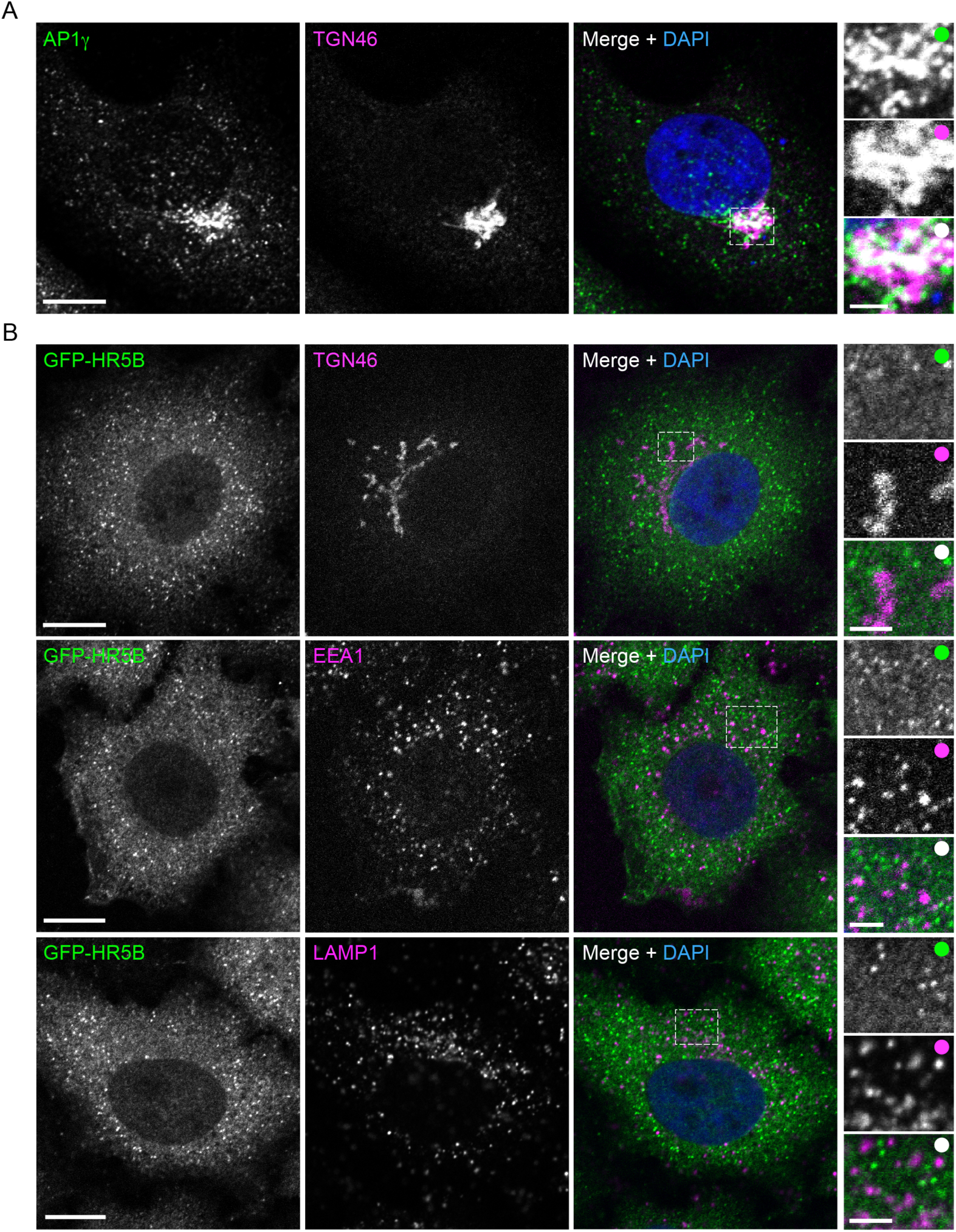
Localisation of AP1γ and GFP-HEATR5B with respect to other structures in fixed human cells. **(A, B)** Representative confocal images of wild-type (A) and stable GFP-HEATR5B (B) HeLa cells stained with antibodies to the indicated proteins (note that GFP signal in panel B was amplified with GFP antibodies). Dashed box shows area magnified in right-hand images. Panel A shows that AP1γ is clustered at the periphery of the TGN. Panel B shows that association of GFP-HEATR5B (HR5B) with TGN46, EEA1 and LAMP1 is rarely observed. Scale bars: main panels, 10 μm; insets, 2.5 μm.

**Figure S3.**
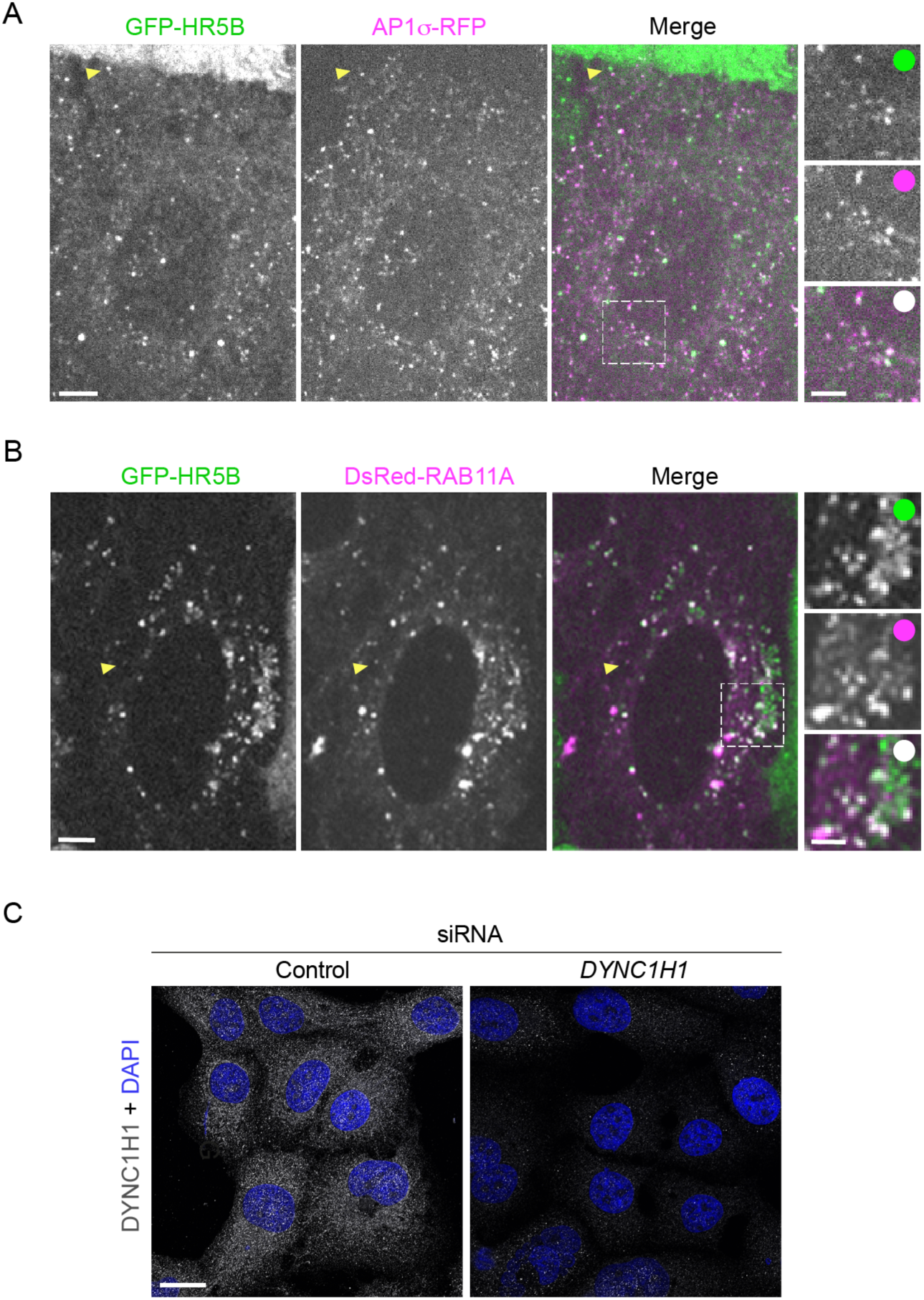
Association of GFP-HEATR5B with AP1γ and RAB11A in live cells and validation of *DYNC1H1* siRNA. **(A, B)** Representative spinning disk confocal images of HeLa cells that have a stable integration of a GFP-HEATR5B (HR5B) construct and have been transfected with AP1σ-RFP or DsRed-RAB11A expression plasmids. Dashed boxes show areas magnified in right-hand images. Arrowheads show particles highlighted in Movies S2 and S4 that undergo long-range transport. **(C)** Representative confocal images of HeLa cells treated with control or *DYNC1H1* siRNAs and stained with a DYNC1H1 antibody, confirming effective knockdown of the protein with *DYNC1H1* siRNAs. Scale bars: A and B, 5 μm; A and B insets, 2.5 μm; C, 20 μm.

**Figure S4.**
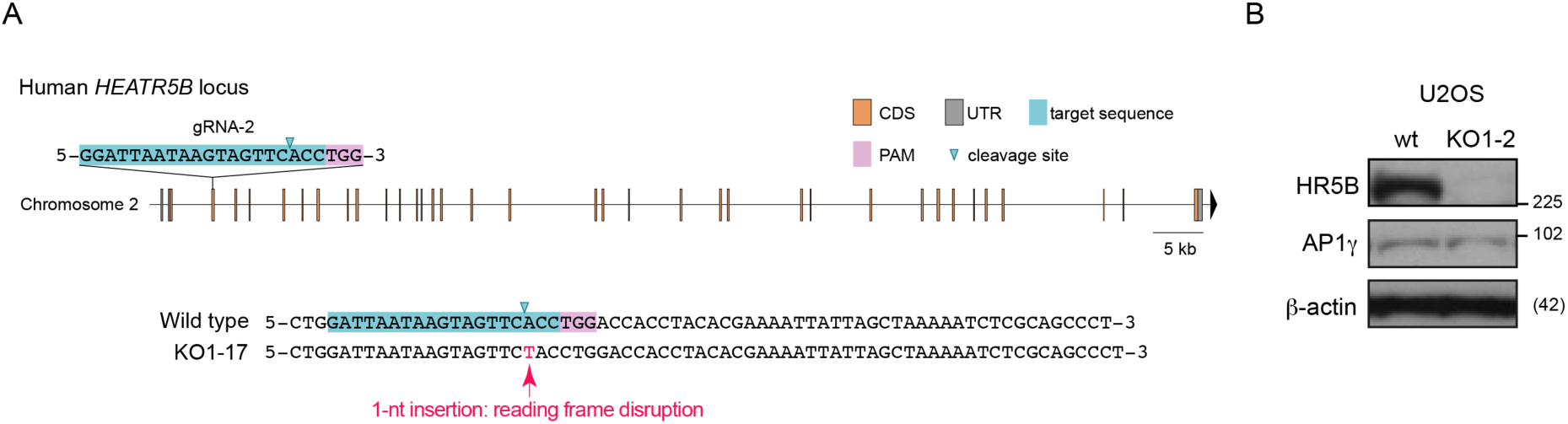
Strategy for generating HEATR5B deficient human cells with CRISPR/Cas9. **(A)** Position of target site of gRNA in human *HEATR5B* locus and sequence of U2OS clone 1-17 (which was used for mutant analysis unless stated otherwise) compared to the wild-type precursor. Information on indels in the other mutant cell lines is available in Table S3. **(B)** Immunoblots showing loss of HEATR5B (HR5B) protein in U2OS KO1-2 clone. Equivalent data for KO1-17 clone is shown in Figure 4D.

**Figure S5.**
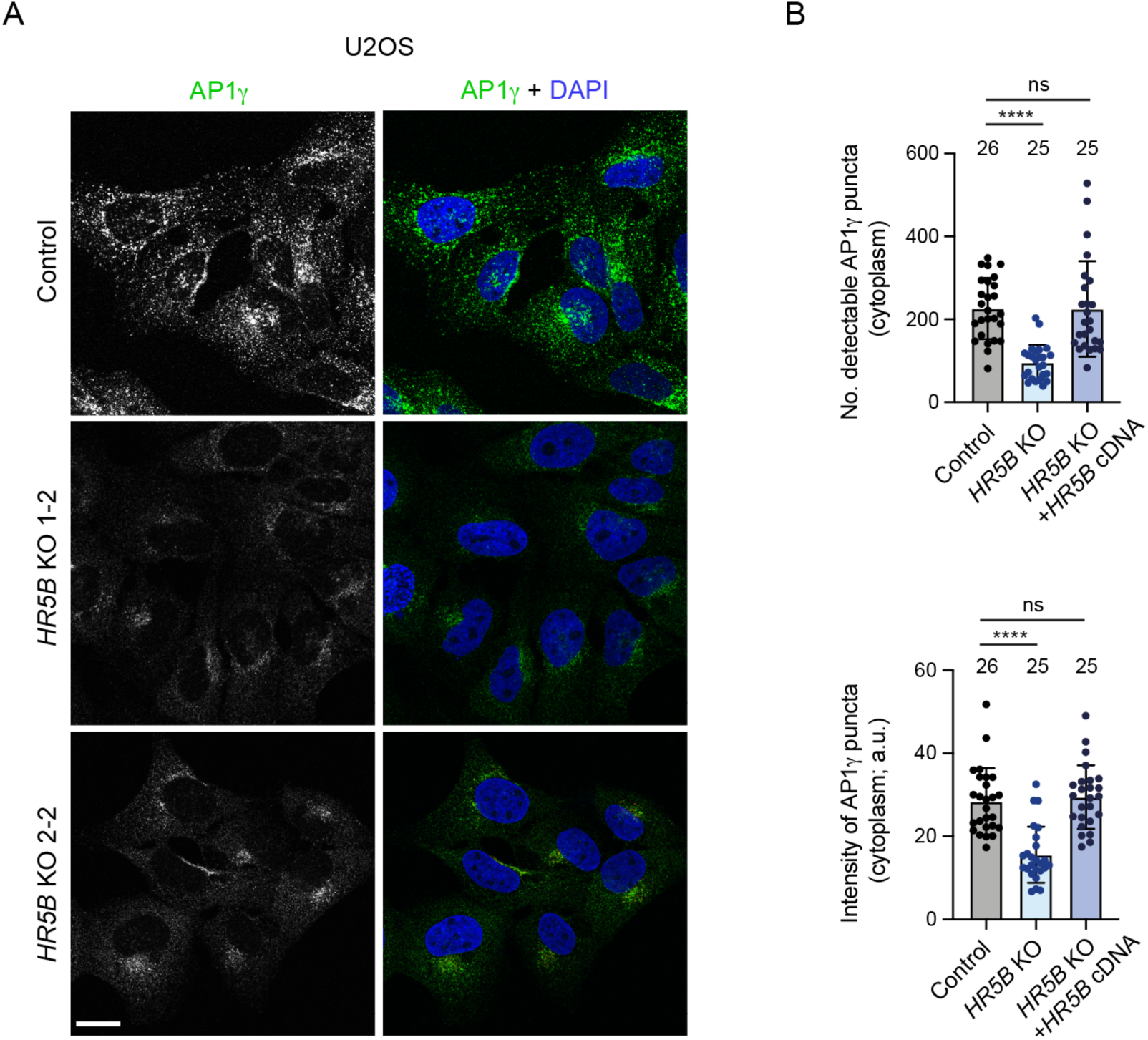
Disruption of AP1γ localisation in additional *HEATR5B* mutant clonal U2OS cell lines and phenotypic rescue with *HEATR5B* cDNA. (A) Representative confocal images of control (parental) and additional *HEATR5B* (*HR5B*)*-*mutant U2OS clonal lines. Scale bar: 15 μm. **(B)** Quantification of number and mean total intensity of AP1γ puncta in control U2OS cells, *HR5B* KO U2OS cells and *HR5B* U2OS cells transfected with a GFP-HR5B expression plasmid (a.u., arbitrary units; see Figure S6 for representative images for rescue condition). Circles indicate values from individual cells, with columns and error bars representing mean ± S.D. Numbers of cells analysed is shown about columns. Statistical significance was evaluated with a one-way ANOVA test. ****, p <0.0001.

**Figure S6.**
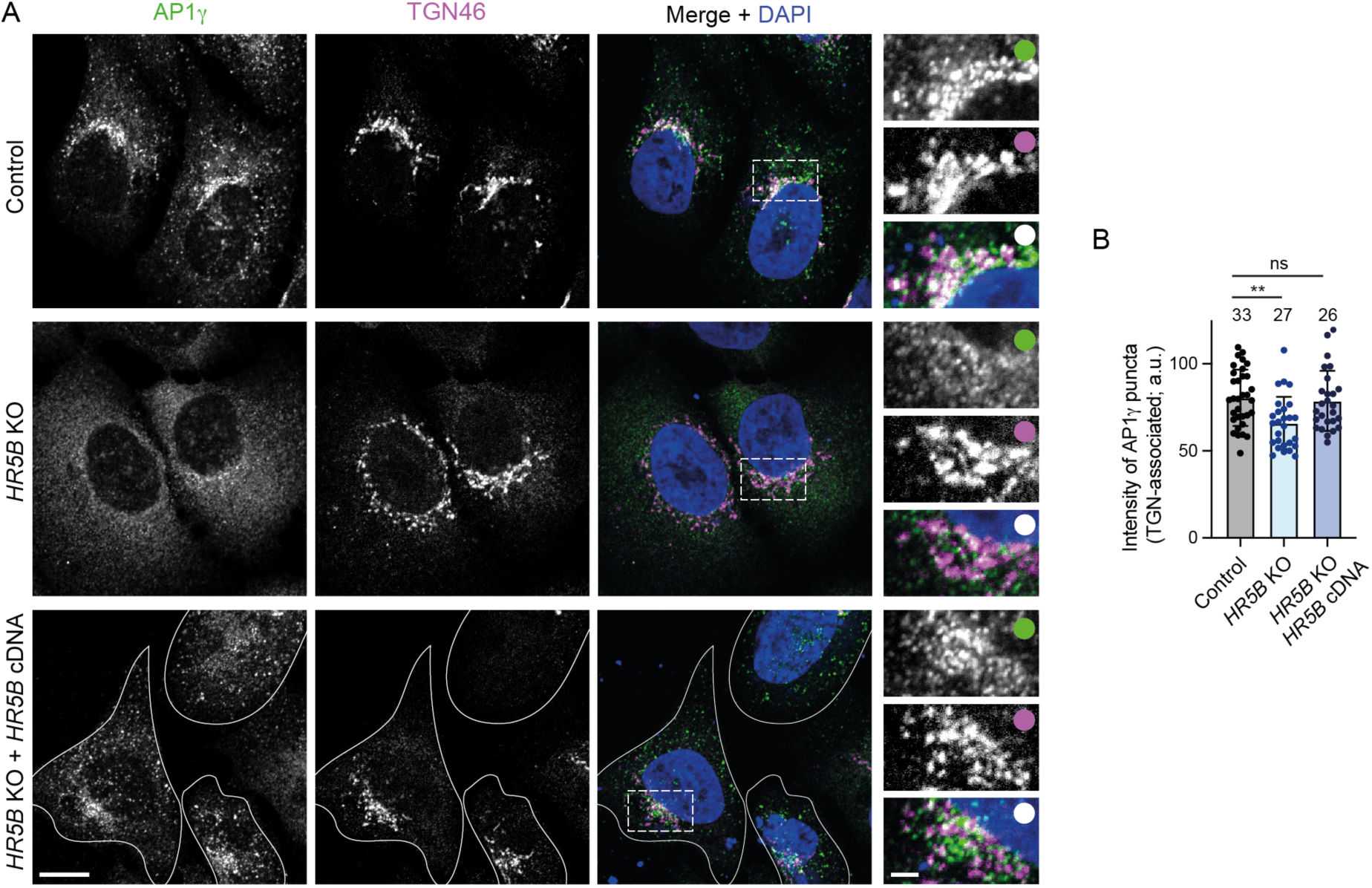
Disruption of HEATR5B partially impairs association of AP1γ with the TGN. **(A)** Representative confocal images of control (parental) U2OS cells, *HR5B* KO U2OS cells and *HR5B* KO U2OS cells transfected with a GFP-HR5B expression plasmid stained with antibodies to AP1γ and TGN46. For illustrative purposes, the *HR5B* KO cells shown are amongst those with a strong reduction in TGN-associated AP1γ signal. Dashed box shows area magnified in right-hand images. White outlines show mutant cells that express GFP-HR5B (as assessed by imaging the GFP channel). Scale bars: A, 10 μm; A insets: 2.5 μm. **(B)** Quantification of mean intensity of TGN-associated AP1γ puncta. Circles indicate values from individual cells, with columns and error bars representing mean ± S.D. Number of cells analysed is shown above columns. Statistical significance was evaluated with a one-way ANOVA test: **, p <0.01.

**Figure S7.**
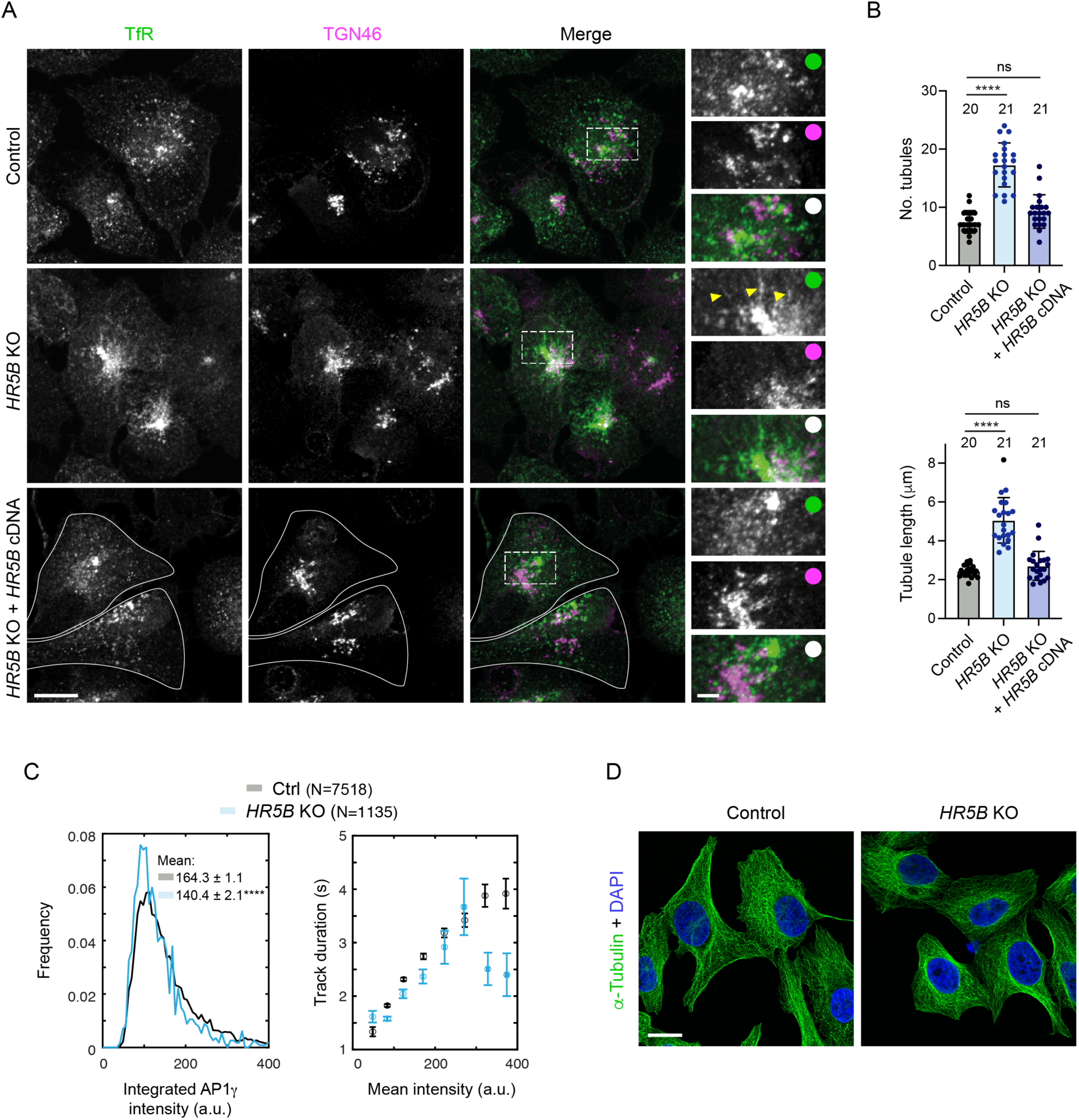
Supplementary data on *HEATR5B* KO phenotypes. **(A)** Representative confocal images of control (parental) U2OS cells, *HR5B* KO U2OS cells and *HR5B* KO U2OS cells transfected with a GFP-HR5B expression plasmid stained with antibodies to Transferrin receptor (TfR), which marks the recycling compartment in association with the TGN, and TGN46. Dashed box shows area magnified in right-hand images. Arrowheads show examples of tubulation in *HR5B* KO cells. White outlines show mutant cells that express GFP-HR5B (as assessed by imaging the GFP channel). **(B)** Quantification of number and length of TfR-positive, TGN-associated tubules. Circles indicate values from individual cells, with columns and error bars representing mean ± S.D. Number of cells analysed is shown above columns. Statistical significance was evaluated with a one-way ANOVA test. **, p <0.01. **(C)** Quantification of AP1σ1-RFP particle intensity (left) and track duration vs mean particle intensity (right) in image series of control and *HR5B* KO live U2OS cells. N = number of particles (from 31 control and 17 KO U2OS cells). In left-hand panel, a shift in the distribution of intensity values to the left in mutant cells shows dimmer fluorescence (statistical significance was evaluated with a Mann-Whitney U test. ****, p <0.0001). **(D)** Representative confocal images of control (parental) and *HR5B* KO U2OS cells stained with α-tubulin antibodies, showing no overt difference in the architecture of the microtubule cytoskeleton. Scale bars: A, 15 μm; A insets, 3 μm; D, 20 μm.

**Figure S8.**
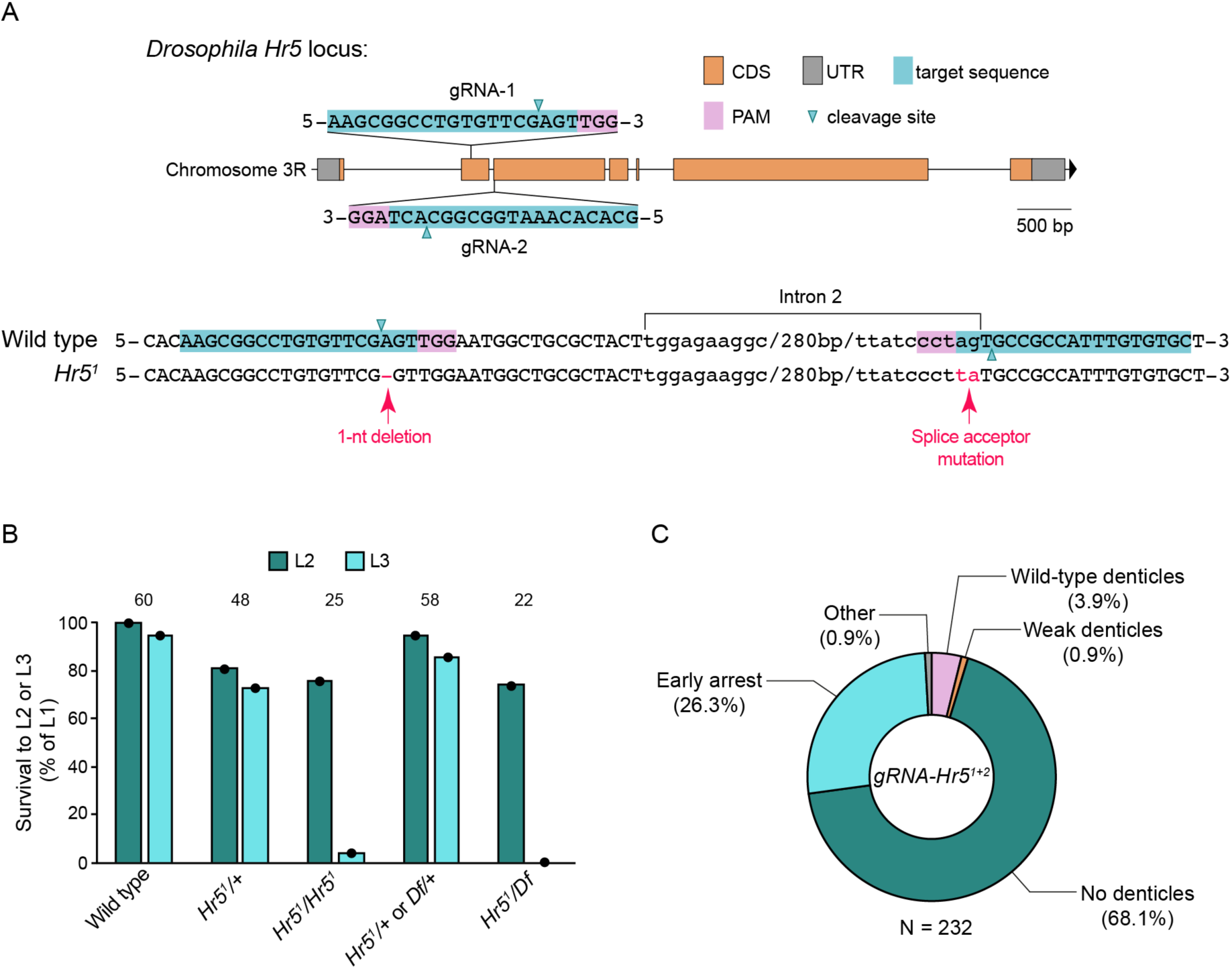
Generation of a *Drosophila Heatr5* mutant allele and analysis of the *Heatr5* zygotic and maternal phenotype. **(A)** Position of target sites of *gRNA-Hr5^1+2^* transgene in *Heatr5 (Hr5)* locus (top) and sequence of the *Hr5^1^* mutant allele compared to the wild-type precursor (bottom). PAM, protospacer adjacent motif. In the sequence alignment, the position of the PAM and target sequence is transposed to the opposite strand for simplicity. **(B)** Lethal phase analysis of *Hr5^1^* zygotic mutants. As none of the genotypes exhibited significant lethality during embryogenesis, only the rate of survival of L1 larvae of the indicated genotypes to L2 or L3 stages was recorded (*Df*: chromosomal deficiency *Df(3R)BSC222*, which uncovers the *Hr5* locus). Data are expressed as a percentage of initial number of L1 larvae for each genotype, with the number of L1 larvae followed for each genotype shown above columns. *Hr5^1^/+* and *Hr5^1^* homozygous larvae were siblings from the same cross, as were ‘*Hr5^1^/+* or *Df/+’* and *Hr5^1^/Df* larvae. **(C)** Quantification of cuticle defects of unhatched embryos from *nos-cas9 gRNA-Hr5^1+2^* females. Embryos in the ‘No denticles’ category had reached late stages of embryogenesis as judged by the development of obvious internal structures, such as abdominal segments and/or mouth parts; ‘Early arrest’ embryos had no obvious internal structures; ‘Other’ represents rare instances of axial patterning defects. Data are pooled from egg lays of females generated in three independent crosses, across which the results were very consistent. N is number of embryos analysed.

**Figure S9.**
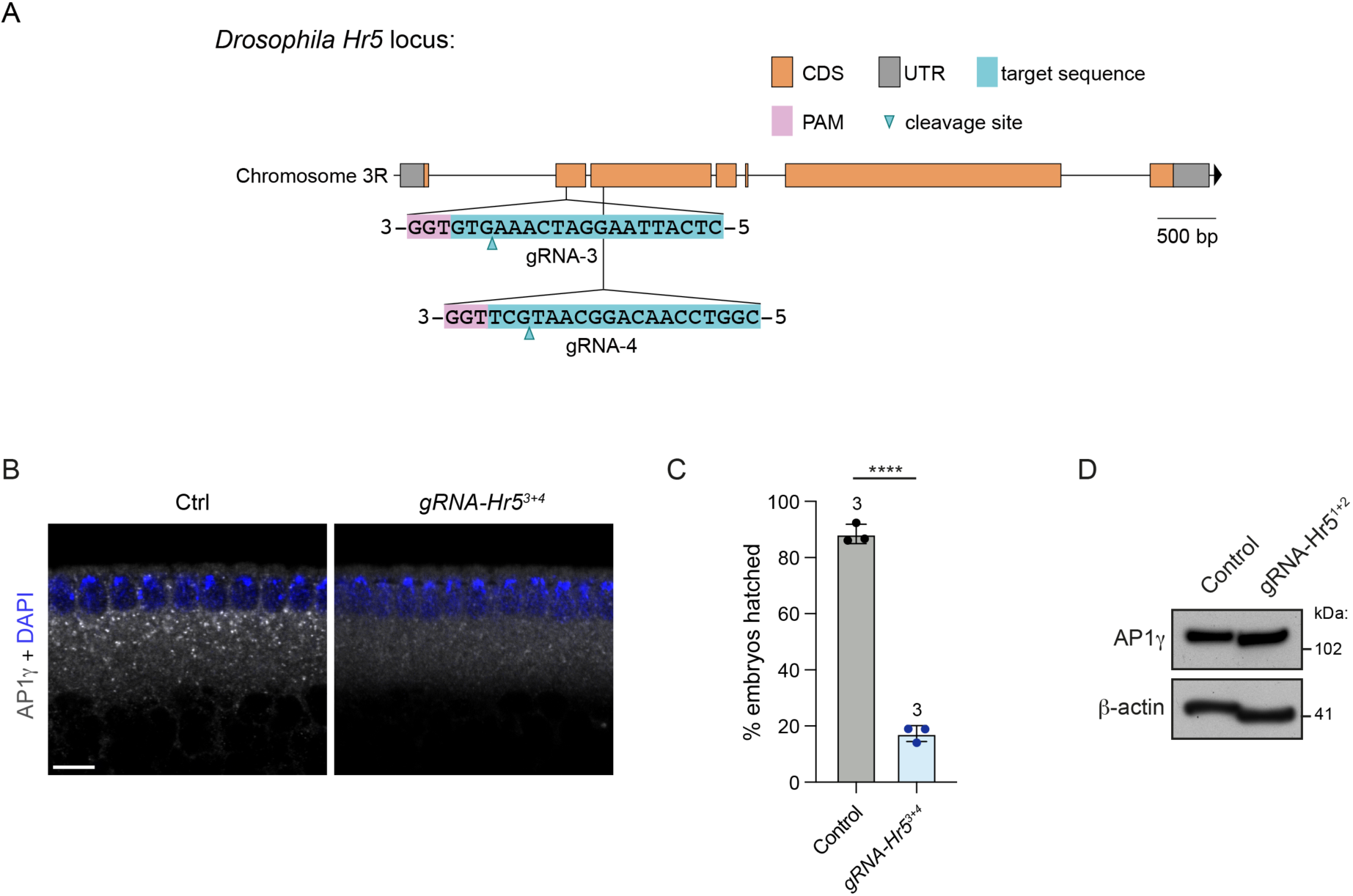
Supplementary results of CRISPR-based depletion of maternally provided Heatr5. (A) Position of target sites of *gRNA-Hr5^3+4^* transgene in the *Heatr5 (Hr5)* locus. **(B)** Representative confocal images of embryos from control (*nos-cas9*) and *nos-cas9 gRNA-Hr5^3+4^* females stained with AP1γ antibodies. Scale bar, 10 μm. **(C)** Hatching frequency of embryos laid by control (*nos-cas9*) and *nos-cas9 gRNA-Hr5^3+4^* females. Chart shows mean values per egg lay ± SD; circles are values for individual egg lays (at least 150 embryos analysed per egg lay); number of egg lays is shown above columns. Statistical significance was evaluated with a t-test: ****, p <0.0001. **(D)** Immunoblot images showing AP1γ protein level in cohorts of 0.5 – 3.5 h embryos laid by control (*nos-cas9*) and *nos-cas9 gRNA-Hr5^1+2^* females. β-actin was used as a loading control.

**Figure S10.**
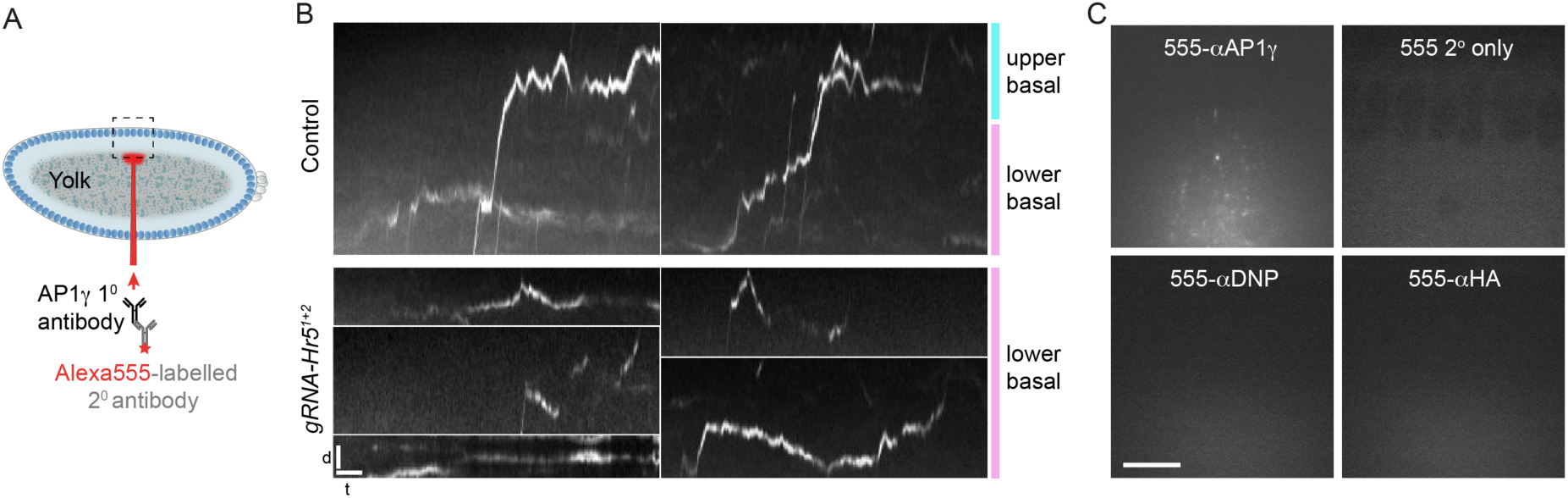
Supplementary information and results for AP1γ motility assays in *Drosophila* embryos. **(A)** Diagram of microinjection procedure. Dashed box shows typical field-of-view for imaging. **(B)** Kymographs showing examples of AP1γ motility in embryos from control (*cas9*) and *nos-cas9 gRNA-Hr5^1+2^* females. Apical is to the top of each image. D = distance and t = time. Scale bars, 2 μm and 20 s. **(C)** Representative images of embryos injected ∼60 s earlier with the indicated antibodies; note the absence of bright puncta for all antibodies except α-AP1γ. Scale bar, 10 μm.

**Figure S11.**
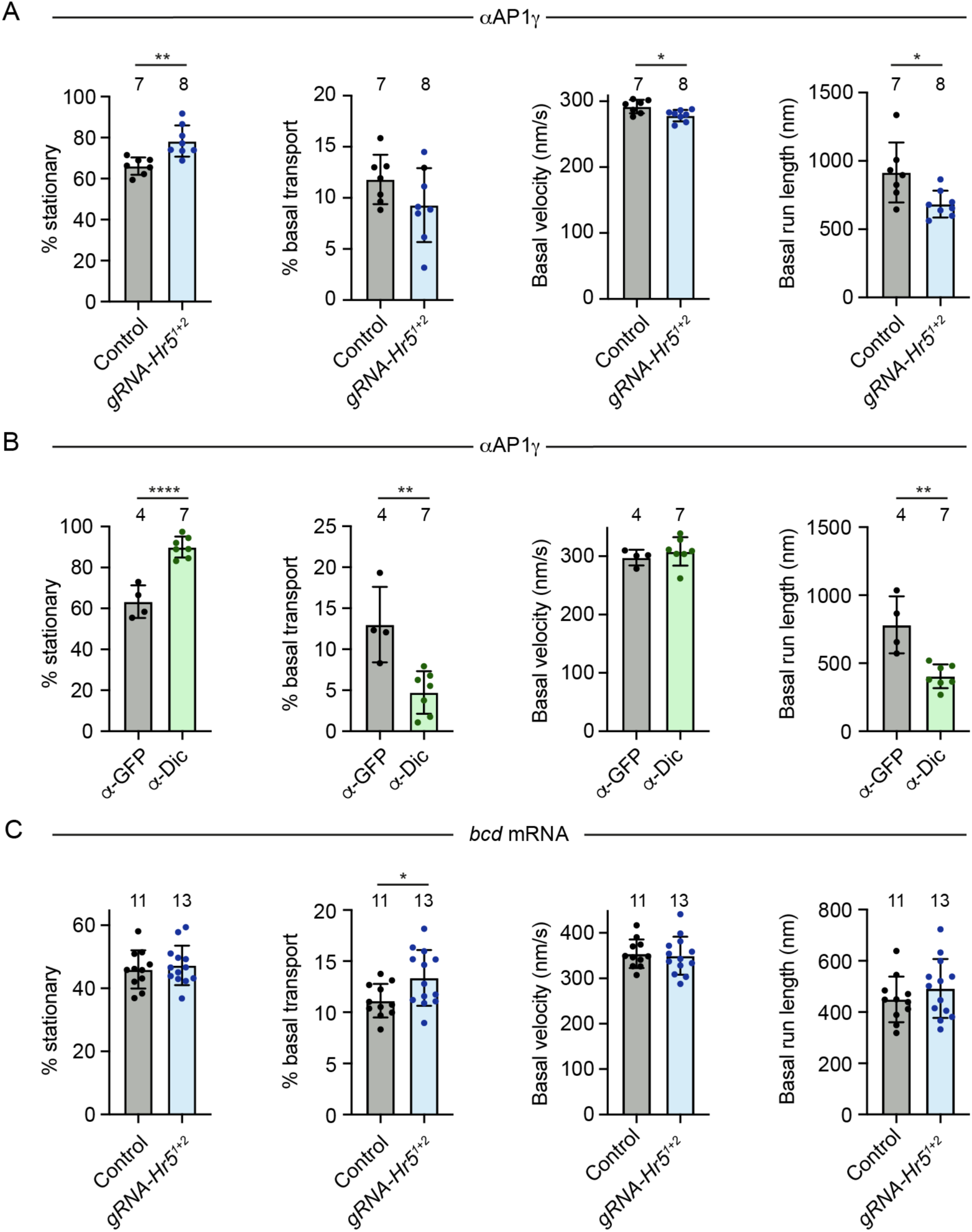
Quantification of frequency of stationary events and basal transport in AP1γ and *bcd* mRNA motility assays. **(A-C)** Quantification of the indicated parameters for AP1γ (A, B) and *bcd* mRNA (C) in embryos of control (*nos-cas9*) and *nos-cas9*, *gRNA-Hr5^1+2^* mothers (A, C) or wild-type embryos pre-injected with function-blocking Dic antibodies or control GFP antibodies (B). ‘% stationary’ and ‘% basal’ are the percentages of particle trajectory time that are classed as immobile or undergoing basal transport, respectively. Circles are mean values for individual embryos; columns and error bars represent means ± S.D. of these mean values; numbers of embryos injected is shown above columns. At least 24 particles were analysed per embryo. Statistical significance was evaluated with a t-test: ****, p < 0.0001; **, p <0.01; *, p <0.05.

**Figure S12.**
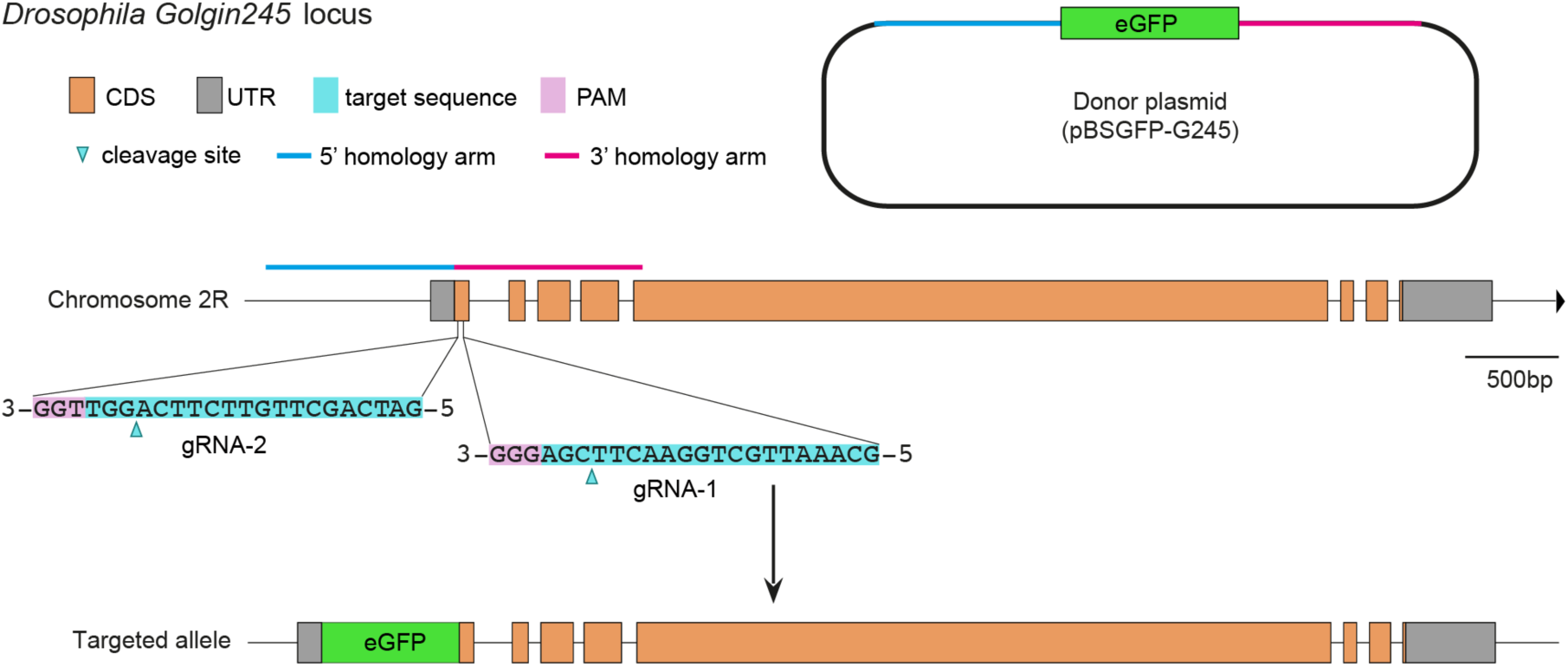
Golgin-245 targeting strategy. Illustration of target sites and homology arms in the *Golgin245* locus, the GFP donor construct, and the *Golgin245* locus after knock-in of GFP.

**Figure S13.**
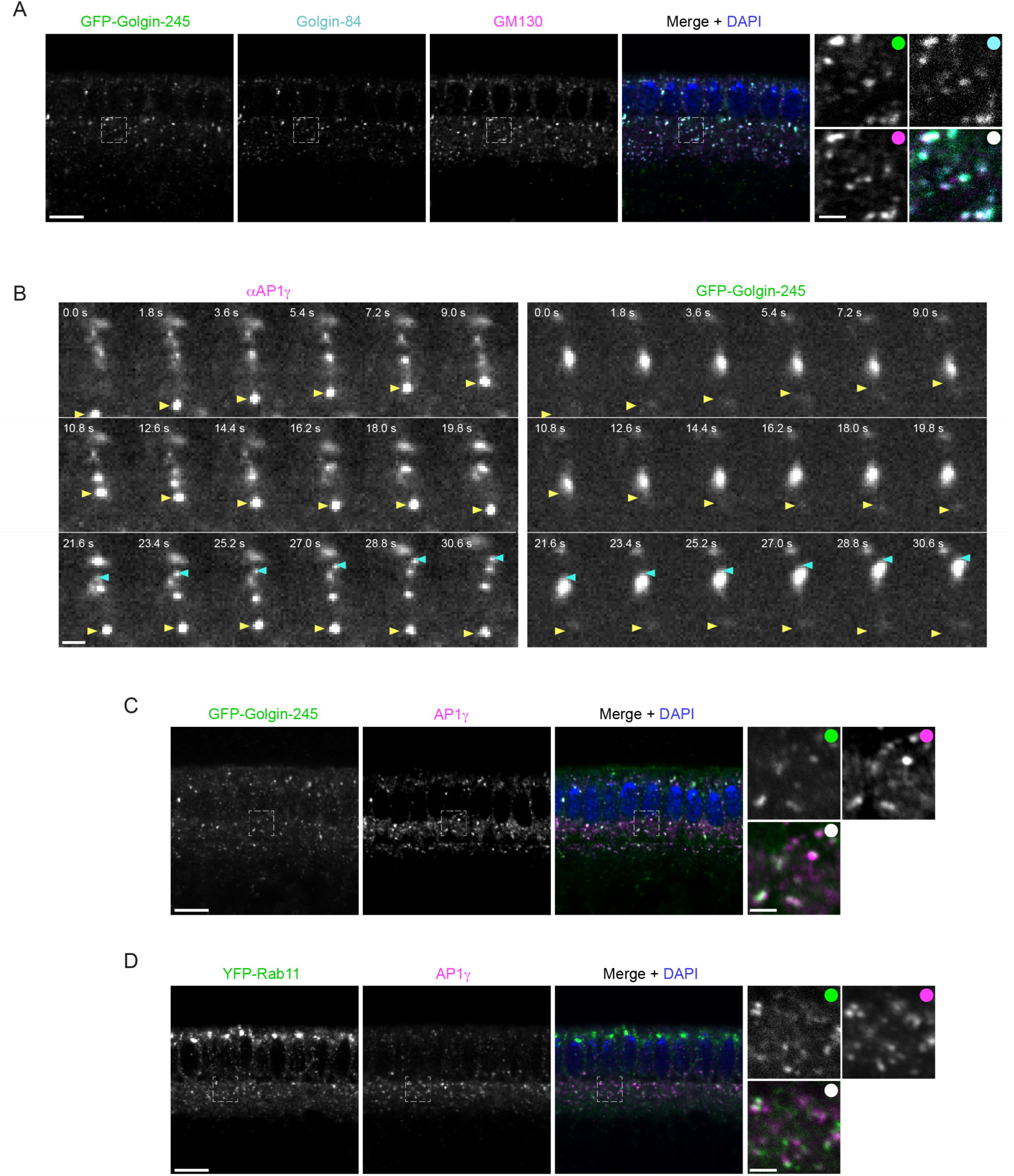
Supplementary information on protein localisation and trafficking in the *Drosophila* embryo. **(A)** Representative confocal images of blastoderm embryo stained for GFP-Golgin-245, Golgin-84 and GM130 showing the GFP fusion protein localises to the Golgi. **(B)** Representative single-channel stills corresponding to the image series from Figure 7A (with yellow and cyan arrowheads in the equivalent positions). **(C, D)** Representative confocal images of wild-type blastoderm embryos stained for GFP-Golgin-245 and AP1γ (C) or YFP-Rab11 and AP1γ (D). Dashed boxes show areas magnified in right-hand images. Scale bars: A, C and D, 10 μm; B, and A, C, D insets, 2 μm.

**Figure S14.**
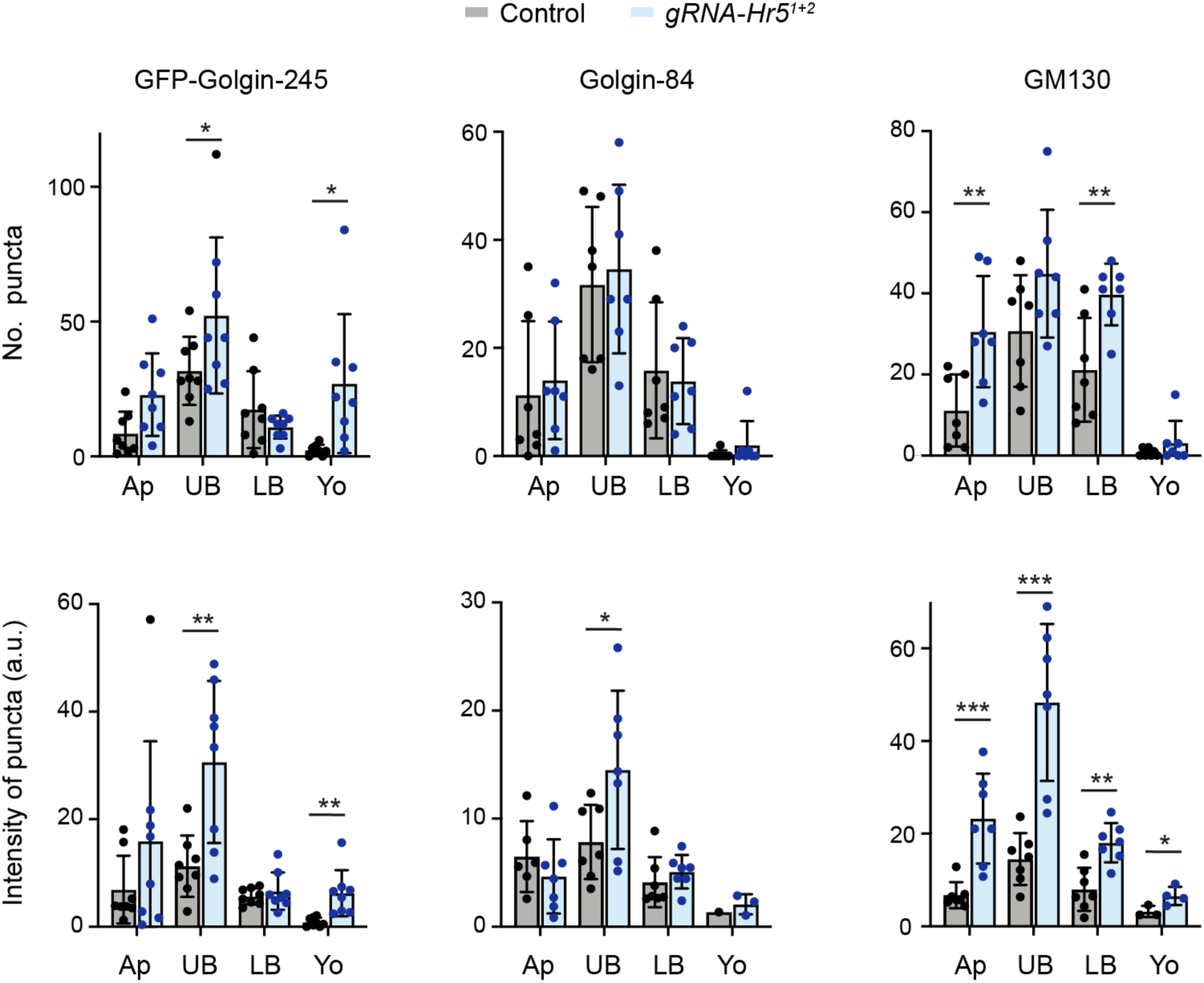
Quantification of Golgi protein localisation in control and *Heatr5* mutant blastoderm embryos. Charts show values for number of puncta and intensity of puncta for the indicated golgin proteins in different regions of the cytoplasm of embryos from control (*nos-cas9*) and *nos-cas9*, *gRNA-Hr5^1+2^* mothers (Ap, apical to the nuclei; UB, upper basal region; LB, lower basal region; Yo, yolk). Columns and error bars show mean of values per embryo ± S.D.; circles show mean values for individual embryos (seven embryos analysed per genotype). At least 50 puncta were analysed per embryo. Statistical significance was evaluated with a t-test: ***, p < 0.001; **, p <0.01; *, p <0.05.

**Table S1.**
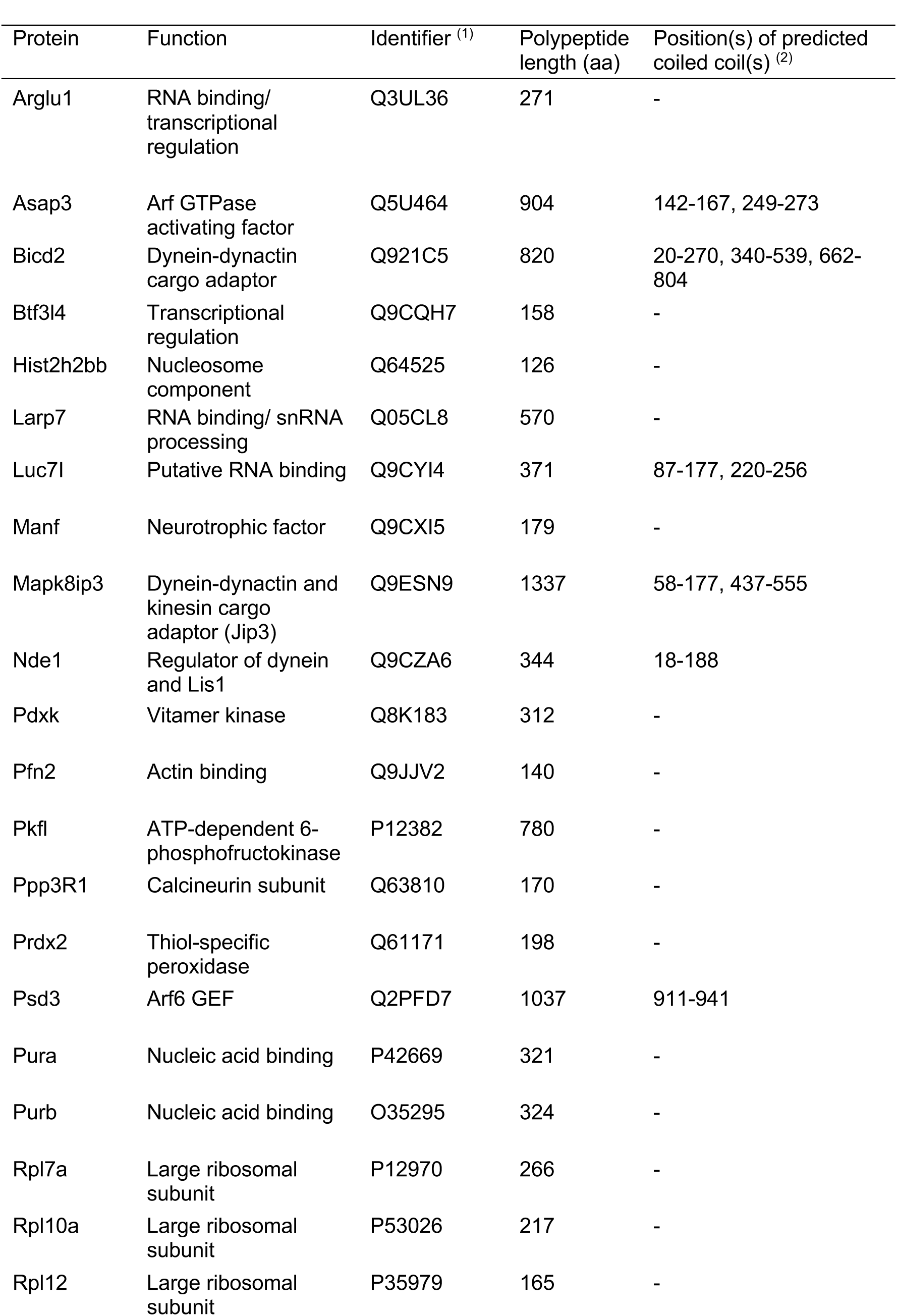
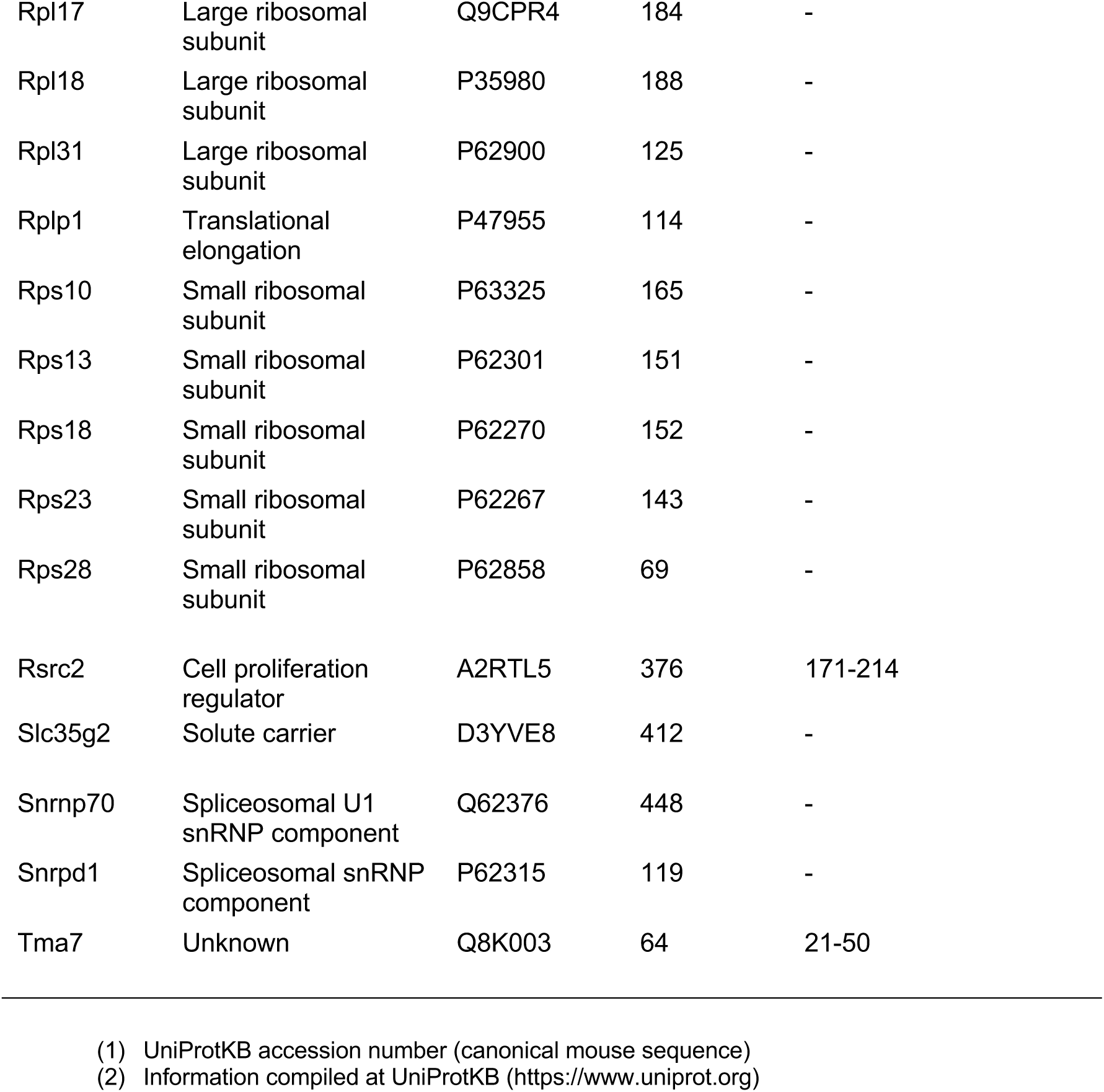
Proteins enriched on the dynein tail vs tag control in the absence of exogenous dynactin in both conditions.

**Table S2.** Proteins enriched on the dynein tail vs tag control in the presence of exogenous dynactin in both conditions.

| Protein | Molecular function | Identifier <sup>(1)</sup> | Polypeptide length (aa) | Position(s) of predicted coiled coil(s) <sup>(2)</sup> |
| --- | --- | --- | --- | --- |
| Acp1 | Phosphotyrosine protein phosphatase | Q9D358 | 158 | - |
| Calm1 | Calcium-based enzyme regulation | P0DP26 | 149 | - |
| Cttnbp2 | Regulates neuronal cortactin distribution | B9EJA2 | 1648 | 119-274 |
| Gfap | Intermediate filament protein | P03995 | 430 | 70-214, 228-374 |
| Heatr5B | AP-1 complex associated | Q8C547 | 2070 | - |
| Hnrnpu | DNA and RNA binding protein | Q8VEK3 | 800 | 626-653 |
| Ppm1e | Protein phosphates | Q80TL0 | 749 | - |
| Rpl14 | Large ribosomal subunit protein | Q9CR57 | 217 | - |
| Rpl19 | Large ribosomal subunit protein | P84099 | 196 | - |
| Rpl6 | Large ribosomal subunit protein | P47911 | 296 | - |
| Rps7 | Small ribosomal subunit protein | P62082 | 194 | - |
| Rps8 | Small ribosomal subunit protein | P62242 | 208 | - |
| Rps16 | Small ribosomal subunit protein | P14131 | 146 | - |
| Smarca2 | Transcriptional regulation | Q6DIC0 | 1577 | - |
| Strip1 | Cytoskeletal regulator | Q8C079 | 837 | - |
| Tial1 | RNA binding | P70318 | 392 | - |
| Wdr91 | Endosomal trafficking | Q7TMQ7 | 748 | - |

(1) UniProtKB accession number (canonical mouse sequence)
(2) Information compiled at UniProtKB (https://www.uniprot.org)

**Table S3.**
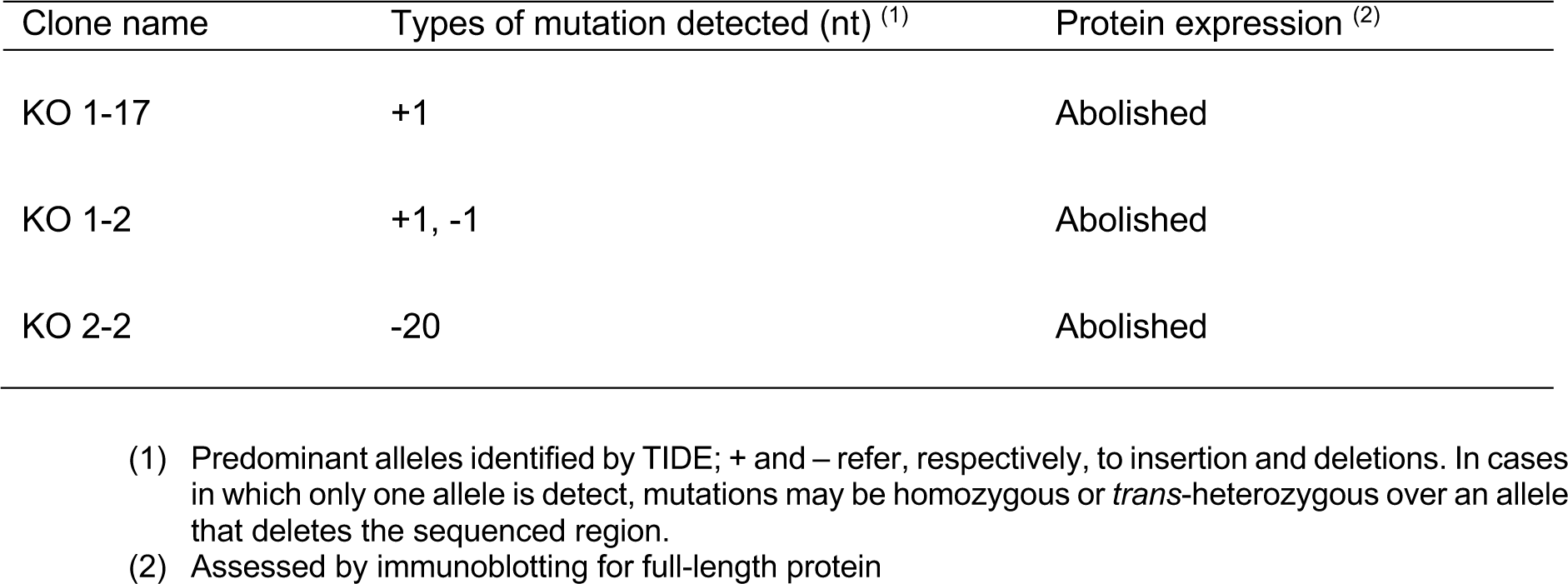
Information on *HEATR5B* mutant U2OS cell lines.

**Table S4.**
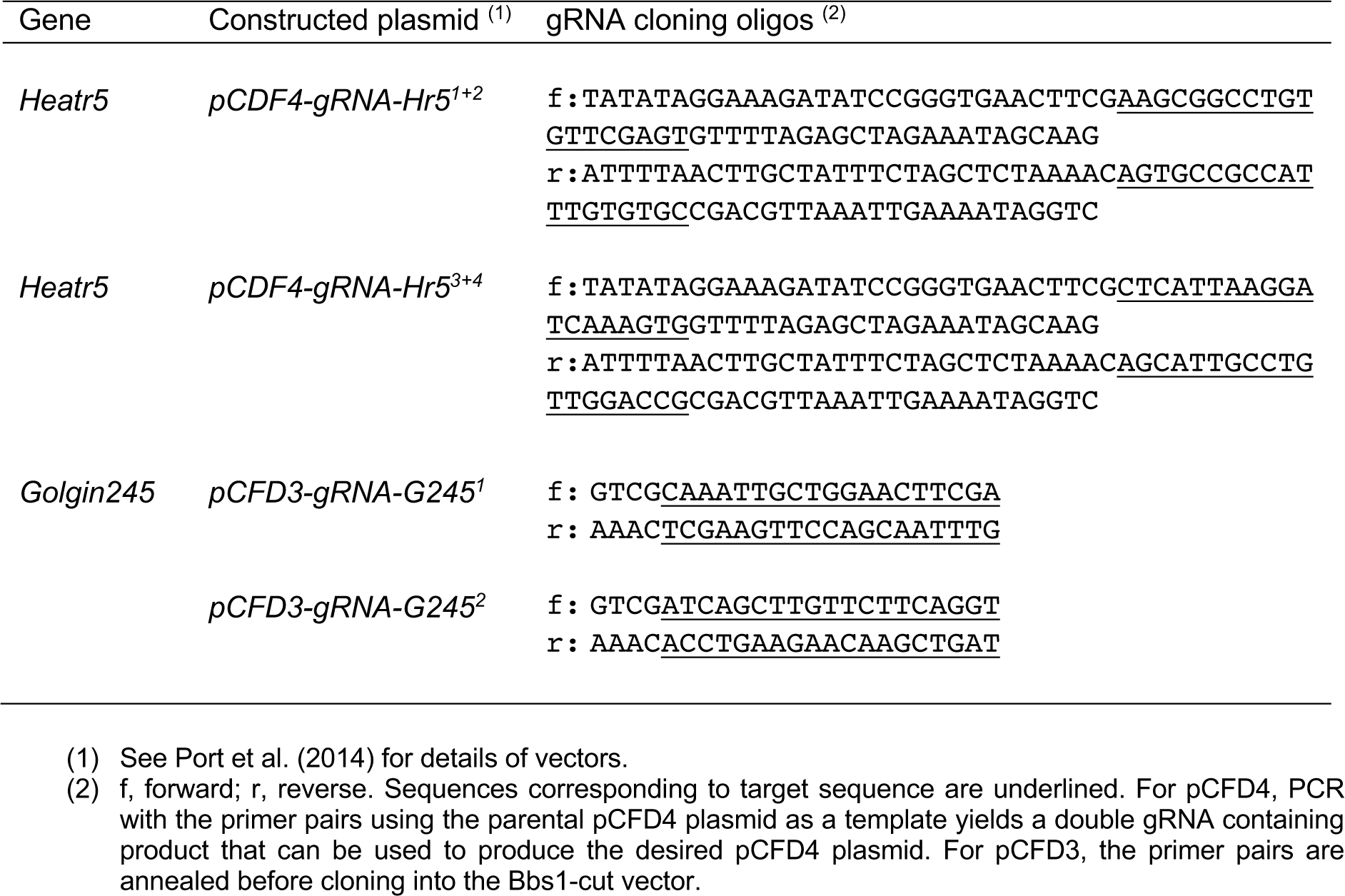
Oligonucleotides used to create gRNA plasmids for CRISPR in *Drosophila*.

## Notes

### Competing Interest Statement

The authors have declared no competing interest.

### Summary of Updates

This is a preliminary revision to address minor comments of reviewers at Review Commons. The revisions includes further discussion of the results in Figure 1, as well as minor changes to improve the clarity of the manuscript.

